# Amyloid aggregates accumulate in melanoma metastasis driving YAP mediated tumor progression

**DOI:** 10.1101/2020.02.10.941906

**Authors:** Vittoria Matafora, Francesco Farris, Umberto Restuccia, Simone Tamburri, Giuseppe Martano, Clara Bernardelli, Federica Pisati, Francesca Casagrande, Luca Lazzari, Silvia Marsoni, Emanuela Bonoldi, Angela Bachi

## Abstract

Melanoma progression is generally associated to increased Yes-associated protein (YAP) mediated transcription. Actually, mechanical signals from the extracellular matrix are sensed by YAP, which activates proliferative genes expression, promoting melanoma progression and drug resistance. Which and how extracellular signals induce mechanotransduction is not completely understood.

Herein, by secretome studies, we revealed an extracellular accumulation of amyloidogenic proteins, i.e. premelanosome protein (PMEL), together with proteins that assist amyloids maturation into fibrils. Indeed, we confirmed the presence of amyloid-like aggregates similar to those detected in Alzheimer disease. These aggregates were enriched in metastatic cell lines as well as in human melanoma biopsies, compared to their primitive counterpart. Mechanistically, we proved that beta-secretase (BACE) regulates the maturation of these aggregates and that its inhibition hampers YAP activity. Moreover, recombinant PMEL fibrils induce per se mechanotransduction promoting YAP activation. Finally, BACE inhibition affects cell proliferation and increases drug sensitivity. These results highlight the importance of amyloids for melanoma survival and the potential of beta-secretase inhibitors as new therapeutic approach to metastatic melanoma.

## Introduction

Melanoma is the most aggressive cutaneous cancer, resulting from the transformation and proliferation of skin melanocytes (Shain & Bastian, 2016). While melanoma only accounts for 1% of skin cancers, it is responsible for the majority of skin cancer deaths with an incidence in rapid increase over the past 30 years. In Europe melanoma accounts for about 150000 new cases, and 27147 deaths/year (Globocan 2018, https://gco.iarc.fr/today/fact-sheets-populations). Approximately 85% of melanomas are diagnosed at early stages when the tumor is thin and surgery is curative in > 95% of cases (American Cancer Society, www.cancer.org). For advanced disease, either unresectable or already metastatic, the therapeutic landscape has benefitted since few years from an unprecedented number of new drugs (e.g. immune checkpoint inhibitors and small-molecule targeted drugs) which have significantly improved the prognosis of advanced patient which is otherwise dismal. That notwithstanding, less than 30% of these cases reaches the 5-yr landmark disease free, clearly indicating that a deeper insight into the biology of melanomas is an unmet clinical need (Guy Jr et al, 2015; Pasquali et al, 2018). The main cause of death is widespread metastases, which commonly develop in regional lymph nodes or in distant organs. Melanoma cells travel along external vessel lattices by regulating adhesion molecules, matrix metalloproteases, chemokines and growth factors. After steadied in the metastatic sites, melanoma cells develop mechanisms that protect them against the attack of the immune system (Zbytek et al, 2008).

Progression to metastatic melanoma is accompanied by increased cell stiffness and the acquisition of higher plasticity by tumor cells, due to their ability to control stiffness in response to diverse adhesion conditions (Weder et al, 2014). During melanoma development, tumor cells are exposed to various types of extracellular matrix (ECM) such as tenascin-C, fibronectin (Frey et al, 2011) and collagen fibers which led to an overall more rigid tumor microenvironment (Yu et al, 2011). Stiffness was suggested to control phenotypic states and to contribute to the acquisition of a malignant phenotype. In epithelial cancers, an extracellular environment characterized by softer matrix enables differentiation, while a stiffer matrix increases proliferation (Lee et al, 2017). Increased ECM rigidity, might also serve as “safe haven” for melanoma cells, protecting them from the effects of chemotherapy; as such, these drug-induced biomechanical niches foster tumor growth and residual disease favoring melanoma resistance (Hirata et al, 2015). It was observed that BRAF inhibitors do not only act on tumor cells but also on the neighboring tumor fibroblasts, paradoxically activating them to produce a stiff, collagen-rich ECM. Melanoma cells fast respond to this new microenvironment by increasing ECM attachment, and reactivating MAPK signaling in a BRAF-independent manner. Understanding how cancer cell-derived ECM is regulated and how it participates in tumor microenvironment remodeling and signaling is critical for developing novel cancer treatment strategies. In this context, secretome studies from tumor and stromal cells provide novel insights in the understanding of the cross-talk between cells within the tumor microenvironment, since they are very sensitive in revealing the key effectors required for the establishment of pre-metastatic niches (Blanco et al, 2012; Kaplan et al, 2005; Karagiannis et al, 2010).

We therefore sought to explore tumor melanoma microenvironment by secretome analysis, investigating the molecular mechanism behind malignant matrix stiffening.

## Results

### *In Vitro* model of metastatic and primitive melanoma

With the aim to understand the functional pathways that differentiate tumor microenvironment of metastatic and primitive phenotype, we investigated two pairs of matched melanoma cell lines. In particular, IGR39 and IGR37 were derived from primitive tumor and lymph node metastasis, respectively, collected from the same 26 years old male patient. Similarly, WM115 and WM266.4 matched cell lines were derived from cutaneous primitive tumor and skin metastasis, respectively, from the same 55 years old female patient (Fig. 1A). Despite their common origin, these cell lines display different phenotypes. In both cases, metastasis-derived cell lines showed a faster growth rate and increased ability to undergo unlimited division compared to the matched primitive tumor-derived cell line (Fig. 1B and Supplementary Fig. S1). Differences in morphology was also denoted between the two matched cell lines: metastasis-derived IGR37 appeared with a short, elongated shape with a spontaneous predisposition to form clusters, while primitive tumor-derived IGR39 remained commonly isolated, displaying higher number of branches and branch elongations (Fig. 1C). On the other hand, IGR39 had higher mobility compared with IGR37 when monitored live using time-lapse microscopy (Fig. 1D, Supplementary Video S1-2). All these data suggest that cells isolated from metastatic tumors grow faster, but move slower than primitive tumors, symptomatic of a proliferative phenotype (Hoek et al, 2008) for IGR37 and an invasive phenotype for IGR39.

**Figure 1.**
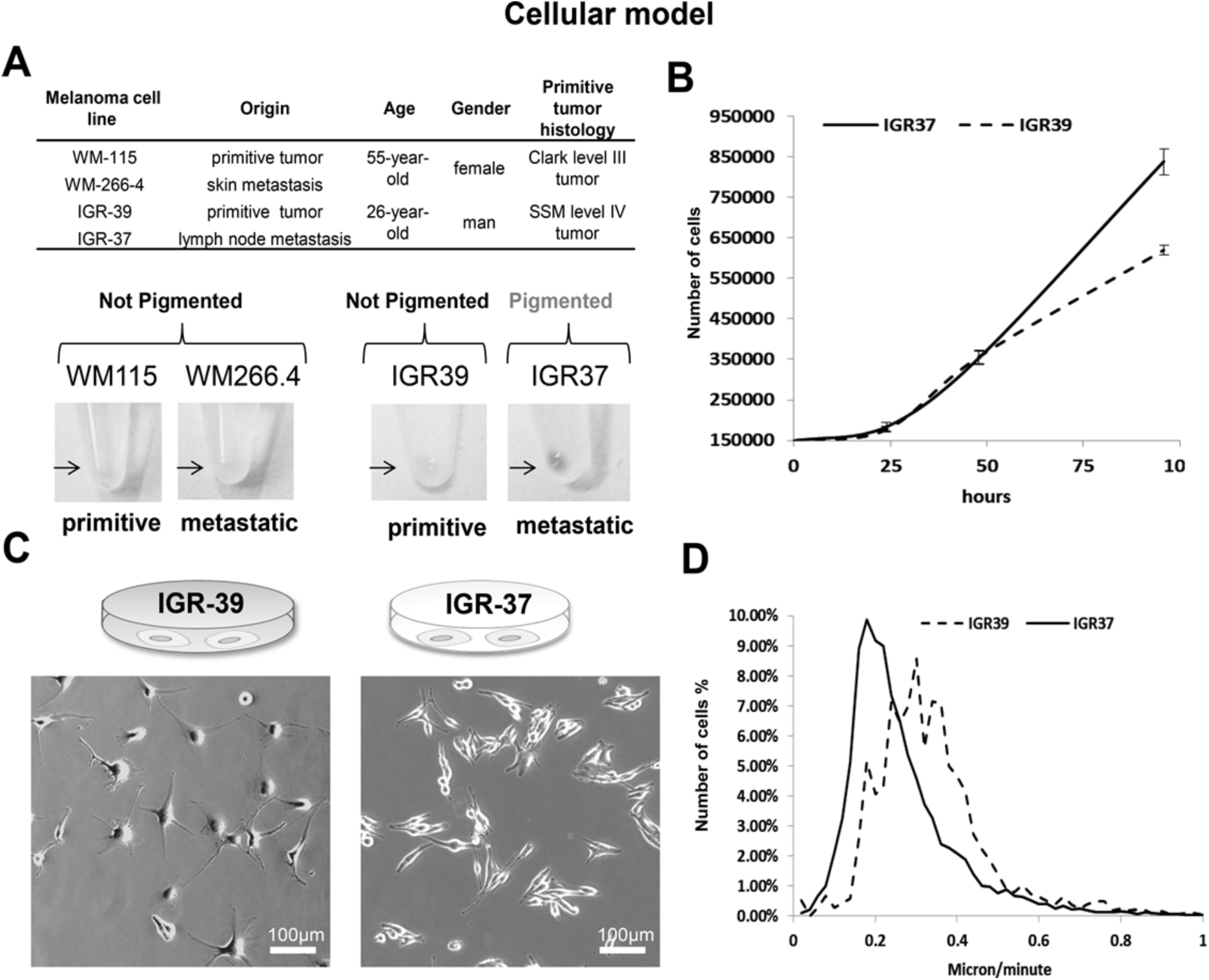
Analysis of a cellular system for primitive and metastatic melanoma. (A) Characteristics of IGR39 /IGR37 and WM115/WM266.4 melanoma cell lines (Upper Panel). Pigmentation of melanoma cells pellets (lower panel). (B) Growth curve of primitive IGR39 and metastatic IGR37 cell line (n=3). (C) Phase-contrast images of IGR39 and IGR37 cells. Scale bar is 100µm. (D) Analysis of IGR39 and IGR37 speed of migration by time-lapse microscopy.

### Global analysis of secreted proteins reveals specific signatures of tumor microenvironment

To investigate the molecular composition of melanoma secretome, we performed a global analysis of the proteins secreted by metastasis-derived (IGR37 and WM266.4) and primitive tumor-derived (IGR39 and WM115) cells lines. To differentiate between proteins that were secreted *versus* the ones present in the serum from cultured conditions, a triple SILAC was performed. We labelled the proteins coming from primitive and metastatic cell line respectively with medium and heavy amino acids (Fig. 2A). For each sample, we analysed the conditioned medium (CM) after 24h of serum deprivation to avoid contamination of high abundant proteins as albumin. We checked for the absence of proteins derived from dead cells by measuring the viability of the cell lines upon starvation and we confirmed that none of them suffered that condition (viability > 95%, Supplementary Fig. S2). As far as secreted proteins are highly glycosylated and this modification might mask proteolytic sites hampering protein digestion, we set up a novel method, named Secret3D (Secretome De-glycosylation Double Digestion protocol), where a de-glycosylation step (PNGase) was added prior to protein digestion performed with double proteolysis to increase protein coverage. Our method enables the unambiguous identification of secreted proteins with high efficiency and quantitative accuracy (Tables S1-S2). As reported in Figure 2A, by examining an equivalent of 500000 cells, we identified 2356 proteins in the SILAC IGR37/IGR39, and 2157 proteins in the SILAC WM266.4/WM115, increasing four times the yield compared to digestion without de-glycosylation (Supplementary Fig. S3A, Peptide Atlas repository) (glycosylated peptides represent about one fourth of the entire dataset, Supplementary Fig. S3B), and improving the proteome coverage if compared to existing methods (Liberato et al, 2018). All proteomics analyses were done in biological duplicate, and for each biological two technical replicates were performed.

**Figure 2.**
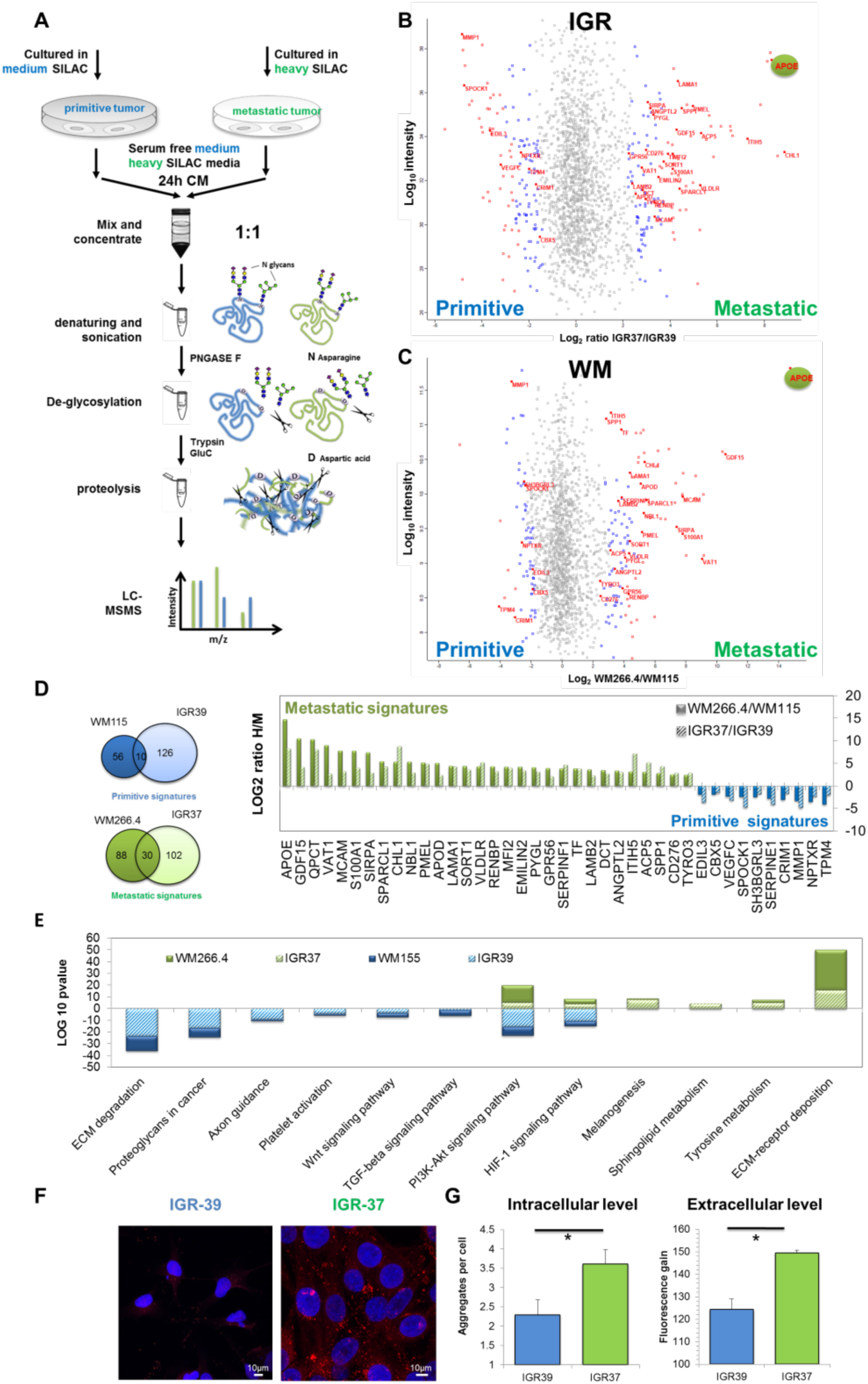
Proteomic analysis of the secretome from primitive and metastatic melanoma cells. (A) MS workflow of Secret3D: Secretome De-glycosylation Double Digestion protocol. (B) Scatter plot of identified and quantified proteins in the secretome of primitive IGR39 and metastatic IGR37. (C) Scatter plot of identified and quantified proteins fin the secretome of primitive WM115 and metastatic WM266.4. (D) Venn diagram of the significant proteins shared by both IGR and WM cell lines. (E) Histograms representing the metastatic and primitive signatures H/M ratios. (E) KEGG Pathway enrichment analysis of the significant proteins. (F) Confocal fluorescence images of Proteostat (1:2000, red spots) and DAPI staining (blue), scale bar is 10µm. (G) Quantitation of aggregates/cell in IGRs cell lines by immunofluorescence analysis, left panel; fluorescence gain of soluble proteins treated with Proteostat reagent, right panel. (T-test analysis, * = p-value<0.05).

By statistical analysis, 270 proteins were found to be differentially secreted in IGR39 versus IGR37. Parallel analysis was conducted in WM115 versus WM 266.4 where the number of differentially secreted protein was 185 (Fig. 2B-C, Supplementary Table S1-2). Results were clustered in order to highlight secretome commonalities. Despite showing only a partial overlap of the differentially secreted proteins (Fig. 2D), both primitive tumor-derived cell lines specifically secrete proteins belonging to ECM matrix degradation (MMP1), Wnt signaling pathway (WNT5a), TGFB signaling pathway (TPM4), proteoglycan degradation (SPOCK1), platelet activation (VEGFC, SERPINE1, EDIL3). These proteins are in agreement with their invasive phenotype, as primitive tumor cells are able to move and invade through the basement membrane or through the vessels walls (Fig. 1D).

Conversely, metastasis-derived cell lines secrete specific proteins belonging to ECM deposition (LAMA1, LAMB2, SPP1), cell adhesion molecule (MCAM, CD276, EMILIN2), lipid transporters (APOE, APOD, PLTP, VLDLR), and melanogenesis related proteins (DCT, KIT1, PMEL) (Fig. 2D-E). Interestingly, APOE is the most secreted proteins in both metastatic cell lines. APOE is a lipoprotein whose primary function is transporting cholesterol, but it also controls the formation of protein aggregates in Alzheimer disease (AD) through the regulation of amyloid-β (Aβ) metabolism, aggregation and deposition. Together with APOE, in metastatic secretome, we found enriched SORT1 and QPCT, proteins known to have a role in facilitating Aβ metabolism (Gunn et al, 2010; Morawski et al, 2014). Notably, in pigmented cells, PMEL maturation is also mediated by APOE and shares similarities with amyloid-β maturation (Van Niel et al, 2015). PMEL amyloid fibrils are known to serve as scaffold for the polymerization of melanin within melanosomes. We found all the above proteins, PMEL included, enriched in the extracellular space highlighting the possibility of fibrils formation extracellularly i.e. plaques.

To test this hypothesis, we used a protein aggregation detection dye (Proteostat). Both pairs of IRGs and WMs matched cell lines were explored: Proteostat staining highlighted an enrichment of aggregated proteins in the metastasis-versus primitive tumor-derived cell line both intracellularly and in the extracellular space (Fig. 2F-G, Supplementary Fig. S1B). To note, amyloid fibrils, together with adhesive proteins, may contribute to the formation of a highly fibrotic extracellular environment specifically in the metastatic cell lines.

### Proliferative and invasive protein signatures are conserved across melanoma cell lines

The secretome signature that distinguishes primitive versus metastatic melanoma was further validated on another cohort of cell lines derived from different patients (Supplementary Fig. S4A). The secretion rate of all the cell lines analysed differs greatly from each other and inversely correlates with growth rate (Supplementary Fig. S4B). These observations were also recently shown by Gregori et al. (Gregori et al, 2014). Analysis was performed using a mixed model based on merged results from dataset normalized either by total number of cells or total protein content. We selected the proteins that were statistically significantly regulated with both approaches. Despite different melanoma cells have different doubling times, different shapes and different proteins secretion rates, we observed highly conserved metastatic and primitive signatures (Supplementary Fig. S4C-D, Supplementary Table S3-4, Peptide Atlas repository). Among the differentially secreted proteins, five were conserved in all the primitive cell lines analysed: MMP1, CBX5, NPTXR, CRIM1 and TPM4; and sixteen proteins were present in all the metastatic cell lines. In accordance with our previous data, APOE, PMEL, QPCT and SORT1 were specifically secreted in all metastatic tumor cell lines together with proteins involved in ECM deposition and adhesive proteins supporting the hypothesis of amyloid fibrils deposition (Supplementary Fig. S4E).

By interrogating Broad-Novartis Cancer Cell Line Encyclopedia (CCLE), proteins involved in amyloid maturation were found enriched in the metastatic cell lines also at the transcriptomic level (Supplementary Fig. S5A). Together with the conserved secretome profile, we found other proteins involved in protein aggregation, such as APP, APLP2 and APLP1, secreted by melanoma in a cell line specific manner. Formation of protein aggregates in the metastatic cell lines was visualized with Proteostat (Supplementary Fig. S5 B).

### APOE and PMEL proteins are overrepresented in the secretome of metastatic melanoma

As discussed before, APOE is the most abundant protein in metastatic secretome. APOE is known to be regulated by liverX receptor (LXR) which is activated by 24- and 25-hydroxycholesterol. In order to verify the involvement of APOE and of cholesterol metabolites in metastatic melanoma, we measured the level of oxysterols in melanoma cells. Indeed, both 24 and 25-hydroxycholesterol were more abundant in metastatic melanoma cells than in primitive tumor cells, thus possibly explaining the proteomic data in matched cell lines (Fig. 3A, Supplementary Fig. S7). Importantly, this data was confirmed also in the other cohort of melanoma cells, where 24-hydroxycholesterol showed the best correlation with APOE levels (Fig. 3B), indicating a specific cholesterol metabolism activation in metastatic melanoma. These evidences sustain the activation of LXR receptor in the regulation of APOE expression in metastatic melanoma, similarly to what reported in astrocytes by Abildayeva and coworkers (Abildayeva et al, 2006), thus enhancing the maturation of PMEL into amyloid fibrils.

**Figure 3.**
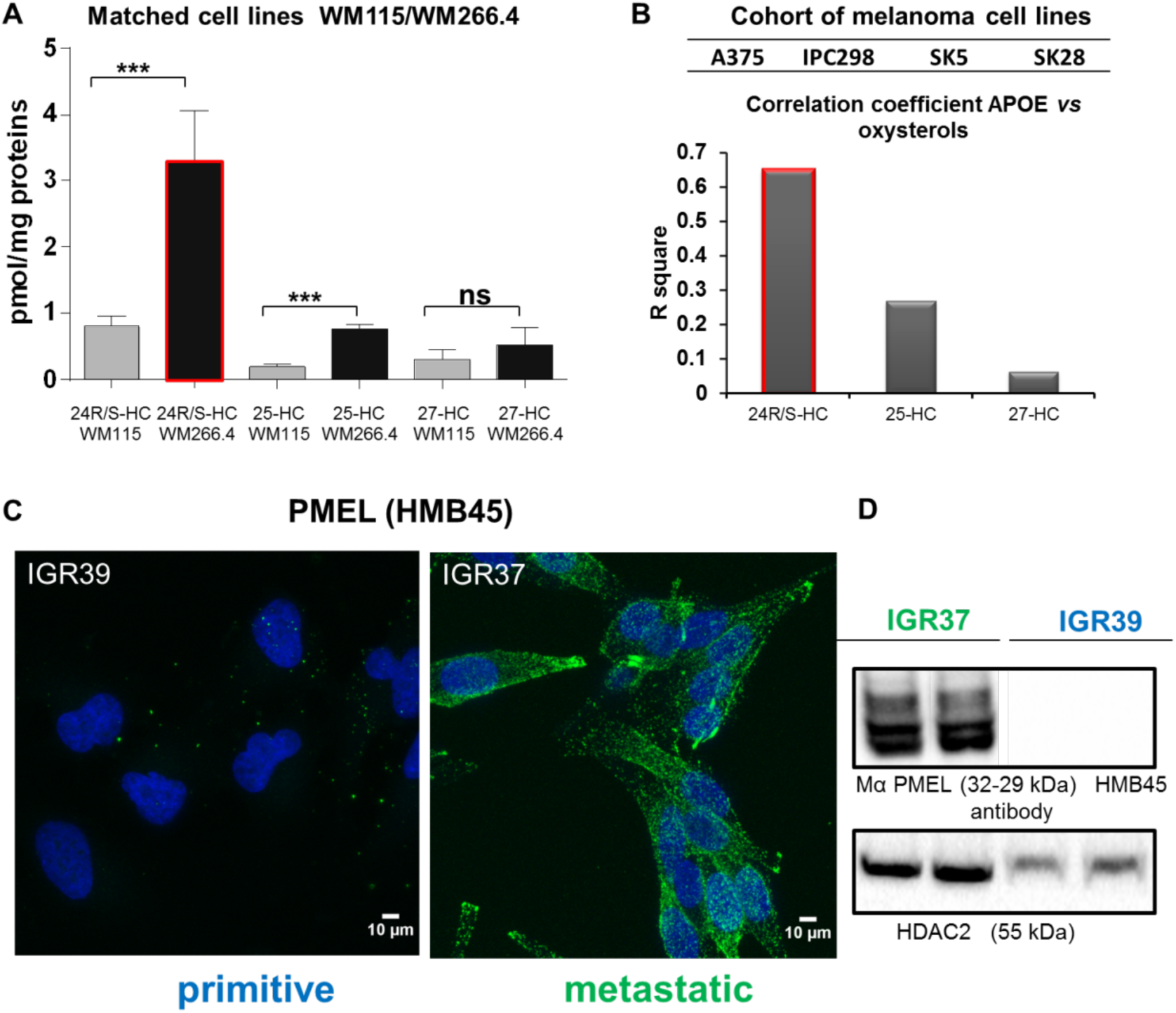
Oxysterols quantification in primitive and metastatic melanoma cells and PMEL expression in IGR37 and IGR39 melanoma cell line. (A) Absolute quantitation of 24, 25 and 27-hydroxycholesterol in primitive and metastatic melanoma cells as indicated. (T-test analysis, ***p-value<0.001). (B). Upper panel: Histogram representing R square of correlation analysis (lower panel) between absolute quantitation of 24, 25 and 27-hydroxycholesterol and label free quantitation of APOE in a cohort of primitive and metastatic melanoma cell lines as indicated. (C) Confocal fluorescence images of anti-HMB45 PMEL antibody signal (green) and DAPI staining (blue) in IGR37 and IGR39. Scale bar is 10µm. (D) Western blot on IGR37 and 39 cellular lysates probed with anti-HMB45 PMEL antibody and anti HDAC2 to check the loading of similar amount of total lysates.

We then verified the presence of PMEL amyloidogenic fragments in metastatic melanoma cells by western blot analysis using an antibody that recognizes the mature form of the protein. As reported in figure 3C, PMEL is exclusively expressed in metastatic cells and not in their primitive counterparts and the molecular weight corresponds to the mature form.

### Aggregated proteins accumulate in metastatic lesions of melanoma patients

Starting from the observation that amyloid–like aggregated proteins were found enriched particularly in the secretome of metastatic melanoma cell lines, we explored if protein aggregates are indeed present in melanoma patients’ tissues. To this aim, we examined samples deriving from primitive tumors and metastases. Archival formalin-fixed paraffin embedded (FFPE) specimens collected from skin primitive tumors and differently localized metastatic sites (i.e. skin, brain and lung), were stained with Proteostat and analyzed by high resolution large scale mosaic/confocal imaging. In both primitive and metastatic tumors samples, we detected a weak or absent signal of protein aggregates localized in the healthy region surrounding tumor tissue (Fig. 4A). Conversely, in primitive tumors, protein aggregates were found spreading along the tissues in small isolated regions inside the tumor area (Fig. 4B). Interestingly, a much higher representation of protein aggregates was detected in metastatic melanoma tissues, without any difference of metastases localization (lung, brain, and subcutaneous skin) (Fig. 4C-D). These data support the hypothesis that progression from primitive to metastatic melanoma is accompanied by increased proteins aggregation.

**Figure 4.**
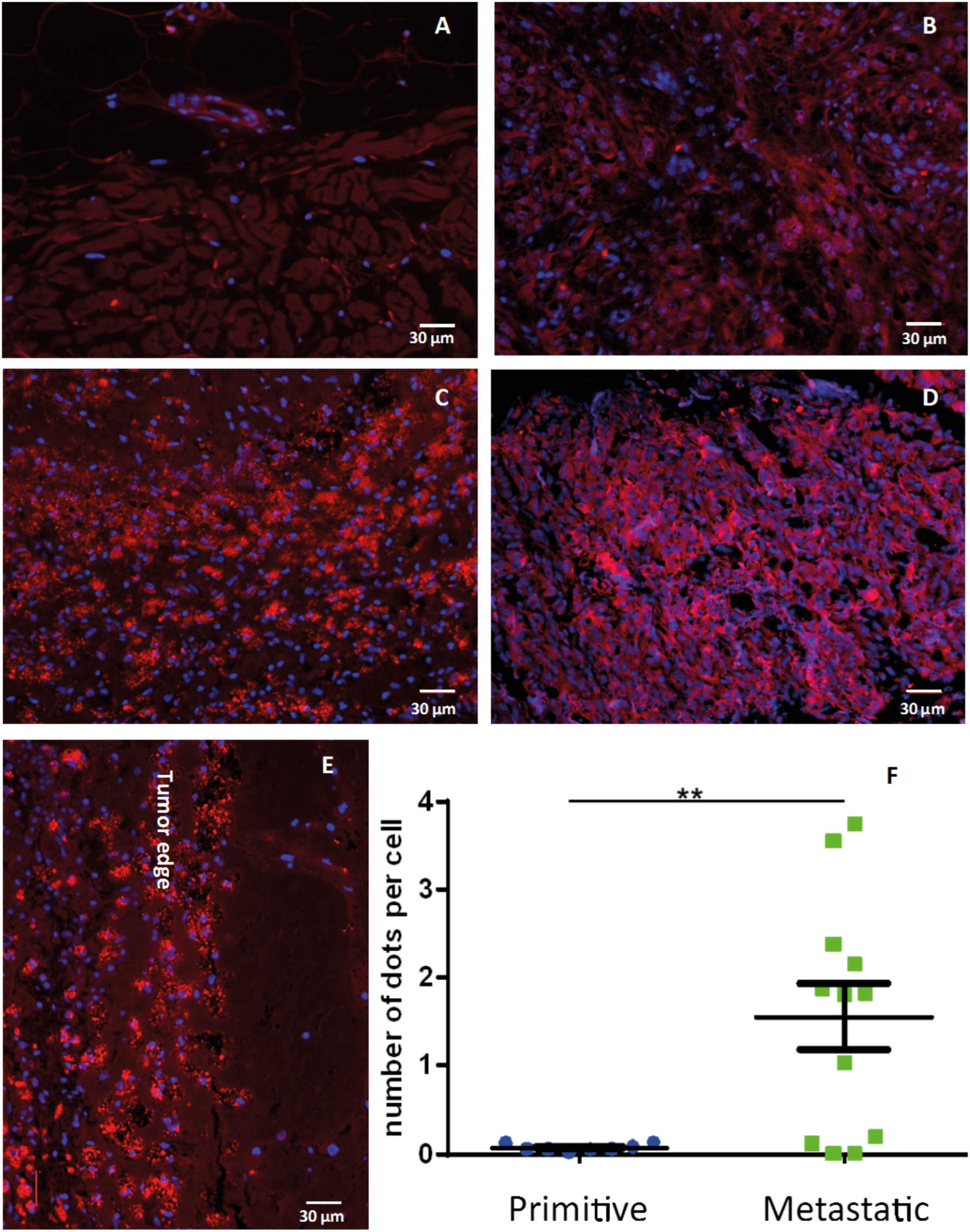
Protein aggregates accumulate in human metastatic melanoma. Immunofluorescence images with Proteostat (red) and DAPI (blue) staining on (A) human normal skin, (B) samples of primitive melanomas, (C) melanoma metastases in brain and (D) melanoma metastases in lung. (E) Details of brain metastases. Scale Bar is 30µm. (F) Quantitation of Proteostat positive dots in primitive vs metastatic melanoma tissues: 6 tissues from metastatic lesions and 6 from primitive melanoma lesions were analysed. For each tissue two sections were quantified. T-test analysis was applied. **p-value= 0.0057.

In details, we observed that protein aggregates appear as dots-like structure on tumor tissue, clearly defining tumor edge (Fig. 4E). Moreover, the number of protein aggregates, quantitated by counting the number of dots per cell, was significantly enriched in metastatic lesions compared to primitive tumors (Fig. 4F). Interestingly, the presence of protein aggregates is not related to pigmentation, as there is no correlation between melanin (hematoxylin-eosin) and Proteostat staining (Supplementary Fig. S6); on the other hand a remarkable correlation with cell proliferation (Ki-67 staining) can be observed (Supplementary Fig. S7), symptomatic of a more proliferative phenotype for the metastatic tissues (Hoek et al, 2008). These results are in accordance with the proliferative phenotype observed in metastatic *versus* primitive cell lines (Fig. 1 and Supplementary Fig. S1).

### PMEL amyloid fibrils drive YAP mediated transcription

After demonstrating the enrichment of protein aggregates in metastatic tissues, we wondered if we could interfere with their production and if this would affect metastasis behaviour. The beta-secretase (BACE 1 and 2) enzymes are known to be involved in the formation of protein amyloids. Indeed, PMEL and APP are cleaved by BACE 2 and 1 respectively, and are able to form mature amyloid fibrils through an APOE mediated process (Rochin et al, 2013). Notably, by interrogating gene expression profiling in TCGA and GTEx dataset, we found that BACE 2 is over-expressed in melanoma more than in any other cancer type (Supplementary Fig. S8A) and correlates with a poor prognosis (Supplementary Fig. S8B). Moreover, melanoma is also characterized by higher mRNA levels of APOE and PMEL with respect to healthy donors (Supplementary Fig. S8C). We therefore choose to pharmacologically inhibit BACE to test if it is actually involved in the formation of the protein aggregates that we observed in melanoma metastasis. NB-360 is a BACE1/2 inhibitor, known to impair the maturation of both APP, in the central nervous system, and PMEL, in normal melanocytes (Neumann et al, 2015). Matched melanoma cell lines i.e. IGRs, were treated with NB-360 at a concentration which is not cytotoxic (Supplementary Fig. S9A-B), but able to decrease the amount of melanin (Supplementary Fig. S9C), indicating an impairment of PMEL amyloidogenic fragments formation (Shimshek et al, 2016). Successively, the secretome was analysed by Secret3D (Supplementary Table S5, Peptide Atlas repository). Notably, primitive and metastatic secretome clustered separately, displaying a different profile which is coherent with the observation of a different phenotype; moreover, NB360 treatment affects both primitive and metastatic cells by targeting different proteins (Fig. 5A). In particular, the amount of secreted PMEL decreased upon treatment in metastatic cells together with other amyloidogenic proteins and known BACE targets (Fig. 5B). The overall impact of the drug was analysed by performing pathway enrichment analysis. Upon treatment, major downregulated pathways were linked to endocytosis, and ECM, together with pathways regulating cell adhesion (Fig. 5C). Among these pathways, we found that the majority of proteins affected by the treatment belong to the metastatic signature identified before(Fig. 5B). Indeed, even if in both primitive and metastatic cells, the same pathways appear to be perturbed, we observed a stronger impact on the metastatic phenotype (Fig. 5C). In particular, confocal microscopy analysis of Proteostat labelled cells after NB360 treatment showed a significant decrease of protein aggregates demonstrating that BACE is involved in their maturation (Fig. 5D-E).

**Figure 5.**
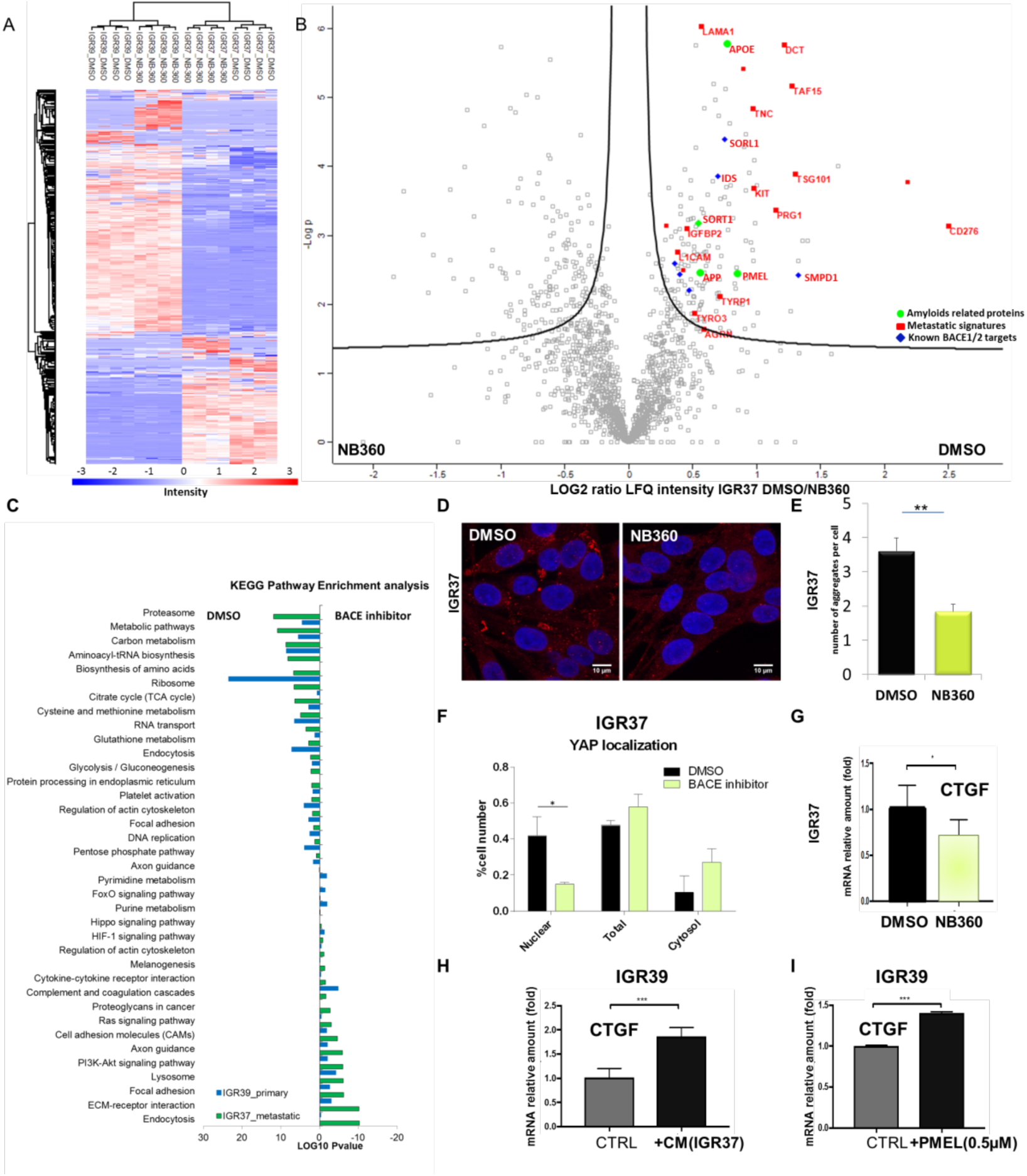
Secretome analysis of IGRs upon BACE inhibition. (A) Unsupervised hierarchical clustering of the proteins identified and quantified in IGR37 and IGR39 upon treatment with DMSO or NB-360. (B) Volcano plot of the proteins secreted by IGR37 cells treated with DMSO or NB-360. (C) KEGG enrichment pathway analysis of the significantly regulated proteins upon BACE inhibition in both IGRs. (D) Confocal fluorescence images of Proteostat signal (1:2000, red spots) and DAPI staining (blue), scale bar is 10µm. (E) Quantitation of protein aggregates in IGRs by immunofluorescence analysis using Fiji software. (F) Quantitation, by immunofluorescence analysis, of YAP in different cellular compartments. Images were quantified by subdividing cells into mostly cytosolic YAP (Cytosol), mostly nuclear YAP (Nuclear), or equal distribution (Total) from three biological replicates. (G) mRNA levels of CTGF measured by real-time PCR in IGR37 treated with DMSO or NB-360. (H) mRNA level of CTGF measured by real-time PCR in IGR39 treated or not with IGR39 conditioned medium (CM), N=4. (I) mRNA level of CTGF measured by real-time PCR in IGR39 supplemented with recombinant PMEL amyloid fibrils (0.5µM), N=3. T-test analysis, * = 0.01<p-value<0.05; ** = 0.001<p-value<0.01; *** p-value <0.001.

Notably, among BACE-downregulated proteins, we identified Agrin (Supplementary Fig. S10A). Agrin is a key protein that senses the extracellular stiffness and activates signaling events to induce the translocation of Yes-associated protein (YAP) into the nucleus (Chakraborty et al, 2017). YAP is a transcription factor that plays an important role in mechanotransduction along with the transcriptional co-activator with PDZ-binding motif (TAZ) (Dupont et al, 2011; Lamar et al, 2012). We postulated that protein aggregates in the extracellular space might increase the external stiffness and activate mechanosignaling leading to YAP mediated transcription. Endorsing our hypothesis, YAP nuclear localization was decreased in response to NB-360 treatment (Fig. 5F, Supplementary Fig. S10B) and YAP target genes, e.g. CTGF, TFGBR2, IGBP4 and FZD1, were found to be downregulated by the drug in the metastatic secretome (Supplementary Fig. S10C). YAP target gene i.e. CTGF is downregulated also at mRNA level upon NB360 treatment, attesting that YAP transcriptional activity is actually impaired by the drug (Fig. 5G).

To further demonstrate that the metastatic microenvironment enriched in amyloidogenic proteins is able to activate a signalling pathway which affects YAP nuclear translocation in a cell autonomous fashion, we supplemented primary melanoma IGR39 cell line with metastatic IGR37 conditioned media and we measured the mRNA level of CTGF as exemplary of YAP target genes. We detected an increased level of CTGF transcription (Fig. 5H), proving that the metastatic secretome is indeed able to modulate YAP. To investigate if the extracellular amyloidogenic proteins act as “mechanotransducers” and are sufficient to activate YAP signalling, we exogenously added recombinant PMEL amyloid fibrils (Fowler et al, 2006) to the primary IGR39 cell line. Interestingly, PMEL fibrils alone are able to increase CTGF expression (Fig. 5I) thus demonstrating that amyloids impinge on a signalling pathway able to activate YAP.

### BACE inhibition impacts on proliferation and enhances chemo-sensitivity in melanoma cells

Convincing evidences are now indicating that ECM confers barriers to treatment as tumor-directed, highly organized ECM structures might inhibit drug penetration and favor cell proliferation (Holle et al, 2016). Therefore, we wondered if NB-360, by changing the ECM organization, might affect metastasis proliferation and chemo-sensitivity. We evaluated the clonogenic activity of IGR melanoma cells upon treatment with NB360, and we found a diminished formation of new colonies (Supplementary Fig. S11A-B) and a decreased proliferation rate (Fig. 6A-B). This effect was similar in both primitive and metastatic cell lines. Conversely, when BACE inhibition is combined with a conventional chemotherapeutic drug such as doxorubicin, metastatic cells become more sensitive to treatment (Fig. 6C-D). Indeed, by evaluating the IC_50_, the combination of BACE inhibition and chemotherapeutic drug resulted in an enhanced synergic effect in metastatic cells while the combinatory effect in primitive melanoma cells appeared to be only additive (Fig. 6E). We can thus speculate that the different response could be related to changes in ECM composition and disappearance of proteins aggregates occurring specifically in metastatic cells as previously demonstrated. Finally, the combined effect was confirmed by a parallel approach where doxorubicin was used at fixed concentration on IGRs and WMs in presence or absence of NB360 (Supplementary Fig. S12A-D). Also in this case, the response to the combinatorial treatment was observed on both cell lines, with a more pronounced effect in metastatic cells.

**Figure 6.**
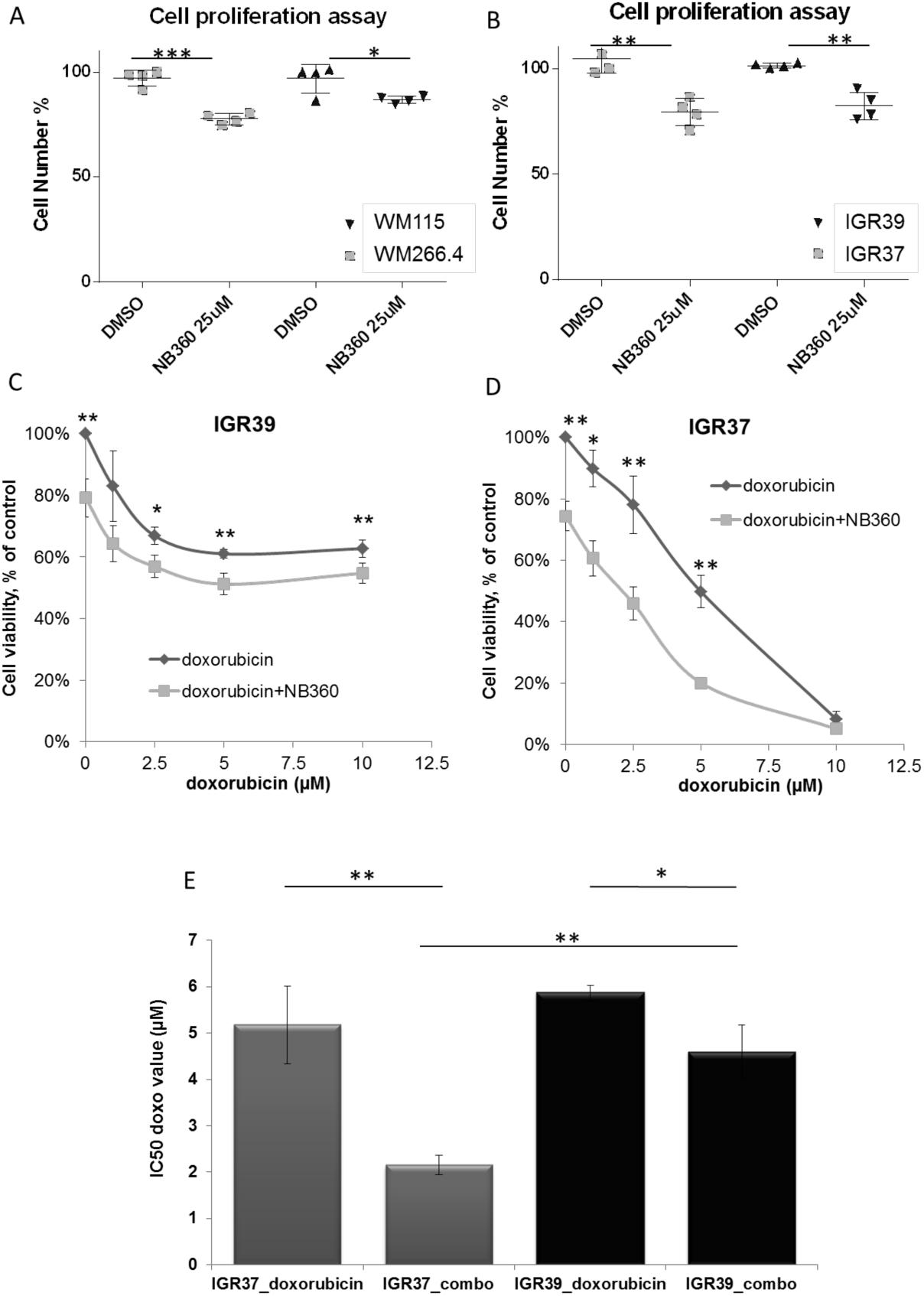
BACE inhibition affects proliferation and chemo-sensitivity in melanoma cells. Panels A-B: MTT assay for IGRs and WMs treated with DMSO or NB-360. Biological replicates N=4. Panels C-D: MTT assay of IGRs treated with NB-360 and different concentration of doxorubicin as indicated. Biological replicates N=4. Panel E: T-test analysis of IC50 values for doxorubicin used alone or in combination with NB-360 (combo). * = 0.01<p-value<0.05; ** = 0.001<p-value<0.01; *** = p-value <0.001.

## Discussion

Cross-talk between tumor cells and its microenvironment has recently gained increasing attention as it actively contributes to cancer progression and metastasis (Wang et al, 2017).

By performing system-level analysis on cellular models of primitive and metastatic phenotypes, we found that protein aggregates were enriched in metastatic cells, both at cellular and extracellular level. Secretome analysis revealed the presence of proteins involved in amyloid deposition enriched in the metastatic microenvironment together with proteins involved in ECM deposition. Altered ECM is frequently observed in various cancers (Lampi & Reinhart-King, 2018) including melanoma (Miskolczi et al, 2018), where stiffening precedes disease development driving its progression through specific mechanical signalling (Pickup et al, 2014). We hypothesize that amyloidogenic proteins in the extracellular space might aggregate, and the deposition of such highly rigid material (Fitzpatrick et al, 2013) might add a further level of stiffness that contributes to activate signalling pathways in melanoma microenvironment. In accordance with our hypothesis, we found APOE as the most secreted protein in the metastatic cell lines. APOE is a lipoprotein, whose primitive function is transporting cholesterol, but it is also involved in the stabilization of amyloid-β fibrils in AD, and of PMEL fibrils in melanocytes maturation (Bissig et al, 2016). APOE expression is regulated by the nuclear LXR activation. In agreement, we showed higher level of the endogenous LXR agonist i.e. 24-hydroxycholesterol in metastatic *versus* primitive cells. LXR activation was observed also in AD (Abildayeva et al, 2006) and, recently, the involvement of oxysterols in tumor progression was reported also in melanoma (Ortiz et al, 2019). In our study, APOE was found associated with the secretion of proteins such as SORT1 (Carlo et al, 2013) and QPCT both involved in amyloid-β fibril stabilization (Morawski et al, 2014). All these observations sustain our hypothesis that in metastatic melanoma extracellular environment there is an overproduction of amyloids like structures. Despite melanoma progression is accompanied by cellular pigmentation (Kirkpatrick et al, 2006; Sarna et al, 2014), we found that also metastatic unpigmented cells, i.e. WM cell lines, actively secrete proteins aggregates.

By analysing tissues from melanoma patients, we highlighted the presence of protein aggregates also *in vivo*. According with our proteomic data, we observed amyloid-like protein aggregation enriched in metastatic lesions compared to primitive tumor tissues. Protein aggregates are hallmark of neurodegenerative disease such as AD, but their involvement in cancer progression is still poorly understood(Xu et al, 2011).

With the aim to interfere with the production of the above mentioned aggregates, we targeted BACE, the enzyme that assists the release of amyloidogenic peptides (Rochin et al, 2013). Interestingly, we found that BACE is highly expressed in melanoma patients compared to healthy donors, and its level of expression correlates with poor prognosis. By using a BACE inhibitor, we reduced the formation of protein aggregates, impaired PMEL and APP shedding, but also affected the secretion of APOE, SORT1 and proteins of the extracellular matrix, as Agrin. In AD, Agrin co-localizes with amyloid plaques and stabilizes amyloid-β fibrils (Cotman et al, 2000); on the other hand, Agrin is also a mechanical sensor that transduces ECM rigidity signals by inducing Yes-associated protein (YAP) activation (Chakraborty et al, 2017). Over the past decade, YAP has emerged as important driver of cancer development (Lamar et al, 2012). In melanoma, YAP has been detected in both benign nevi and metastatic tumor and it was postulated to contribute to both invasive and metastatic behavior (Nallet-Staub et al, 2014). These evidences have encouraged researcher to target YAP activity for anti-cancer therapy (Johnson & Halder, 2014). Here, we showed that BACE inhibition affects YAP nuclear localization and YAP targets expression. Mechanistically, we propose a model where protein aggregates rigidity induces mechanotransduction leading to YAP activation (Fig. 7). Indeed, we demonstrated that metastatic melanoma microenvironment is able to induce YAP mediated CTGF transcription in a cell autonomous way and that PMEL amyloidogenic fibrils extracellularly is able to recapitulate this scenario. We can therefore conclude that, in metastatic melanoma, BACE activity assists the secretion of protein aggregates into the extracellular space, which are sufficient to activate YAP mediated transcription. In this context, we suppose that agrin might participate to this novel YAP signalling cascade, however additional experiments are needed to address this pathway.

**Figure 7:**
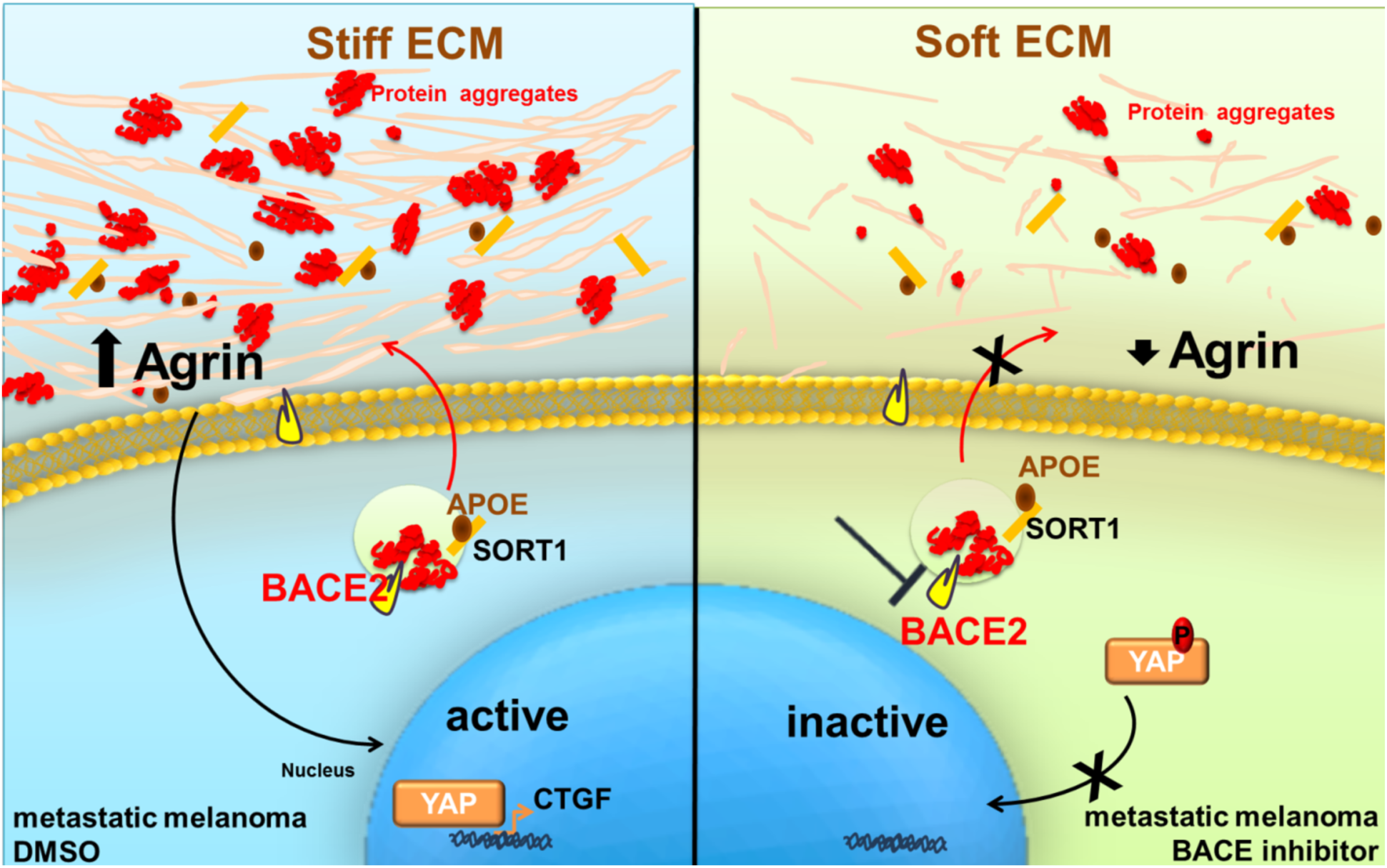
BACE as a new regulator of YAP in metastatic melanoma cells. In metastatic melanoma, BACE activity assists the secretion of protein aggregates into the extracellular space. The presence of these aggregates might be sensed by agrin, known to activate the YAP signalling cascade, and is able to induce YAP mediated CTGF transcription. In turn, melanoma cells treated with BACE inhibitor produce fewer protein aggregates and show a lower YAP transcriptional activity.

In melanoma, YAP overexpression confers resistance to BRAF inhibitor, whereas YAP depletion increases drug sensitivity (Kim et al, 2016). Consistently, we observed that melanoma cells treated with BACE inhibitor become less proliferative and more sensitive to chemotherapy. This combined treatment is more effective on the metastatic cells vs the primary tumor. Indeed, protein aggregates rigidity might contribute in the formation of the “safe heaven”, favoring tumor growth and melanoma resistance (Hirata et al, 2015). Supporting this hypothesis, rigid tumor microenvironment was often associated to the formation of a physical barrier affecting drug uptake (Holle et al, 2016).

Melanoma recent therapies are remarkably efficient in a subpopulation of patients; for those who do not respond though, melanoma remains a devastating disease raising the need of alternative therapies. Here we found a potential new druggable target, i.e. BACE, able to affect melanoma microenvironment. Targeting BACE in combination with chemotherapy might open new revenues to counteract metastatic melanoma. Moreover, this therapy might also interfere with the mechanosignaling pathway that can promote metastatic growth and survival (Lamar et al, 2012). In our work, we have underlined a cell autonomous effect of protein aggregates deposition, but it would be interesting to explore the effect of the presence of amyloid like structures also on neighboring cells. Dr Richard Hynes, from MIT, recently demonstrated in a PNAS paper that in vivo ECM production is mostly fibroblastic, while ECM remodeling is both tumor cell and fibroblastic cell dependent. Here, we provided evidence that amyloid plaques are melanoma cell secreted, therefore they might have an additional contribution to ECM remodelling to the fibroblastic one. Moreover, as amyloidogenic proteins overexpression has been reported also in other tumor types, such as breast(Danish Rizvi et al, 2018) and pancreas(Westermark et al, 2017), it is attractive to think that the same mechanism that we described could be exploited also in other diseases.

## Materials and Methods

### Reagents and tools table

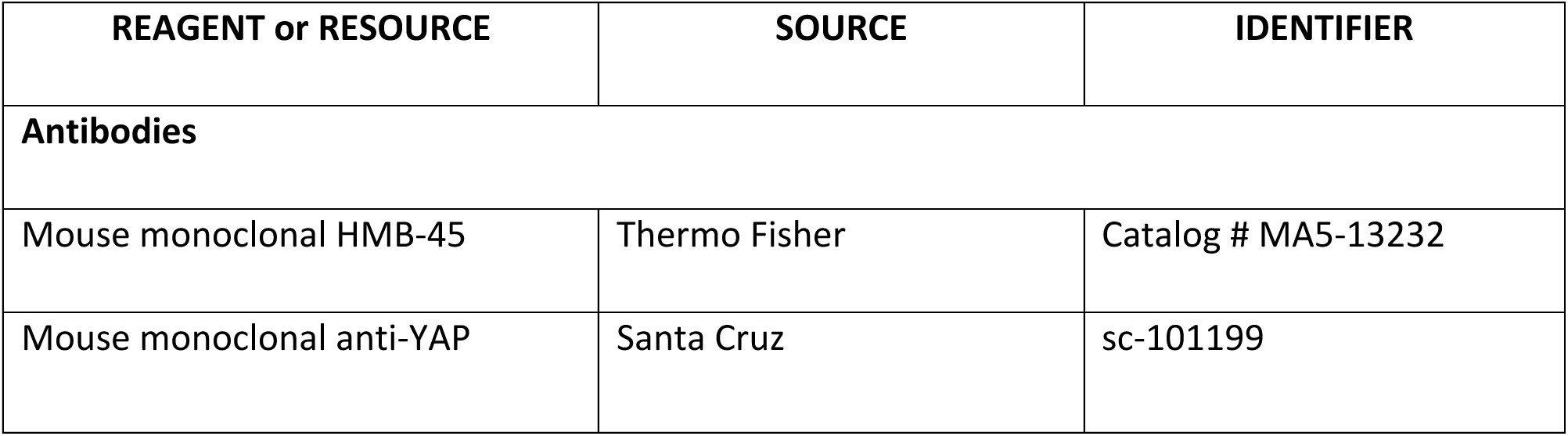
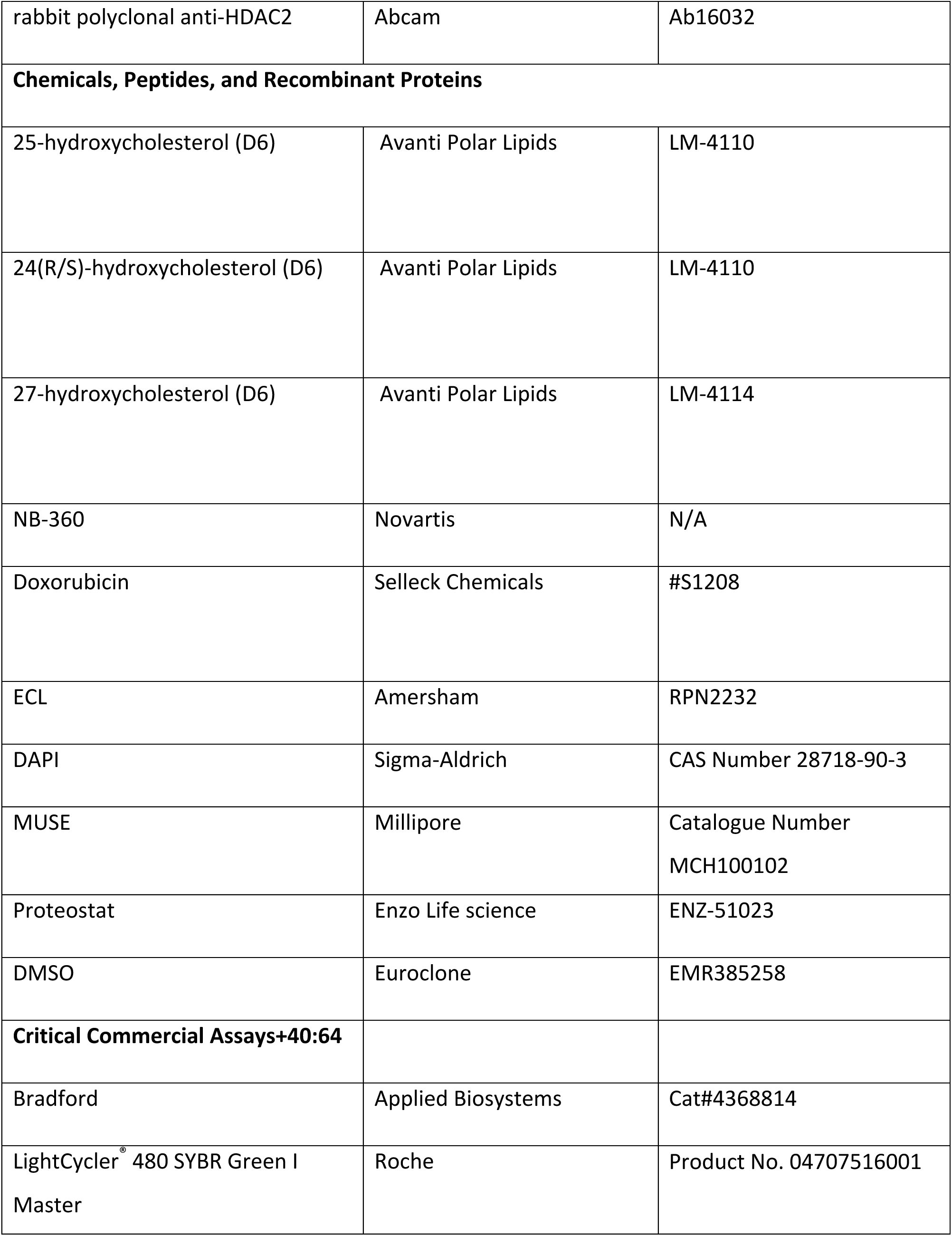
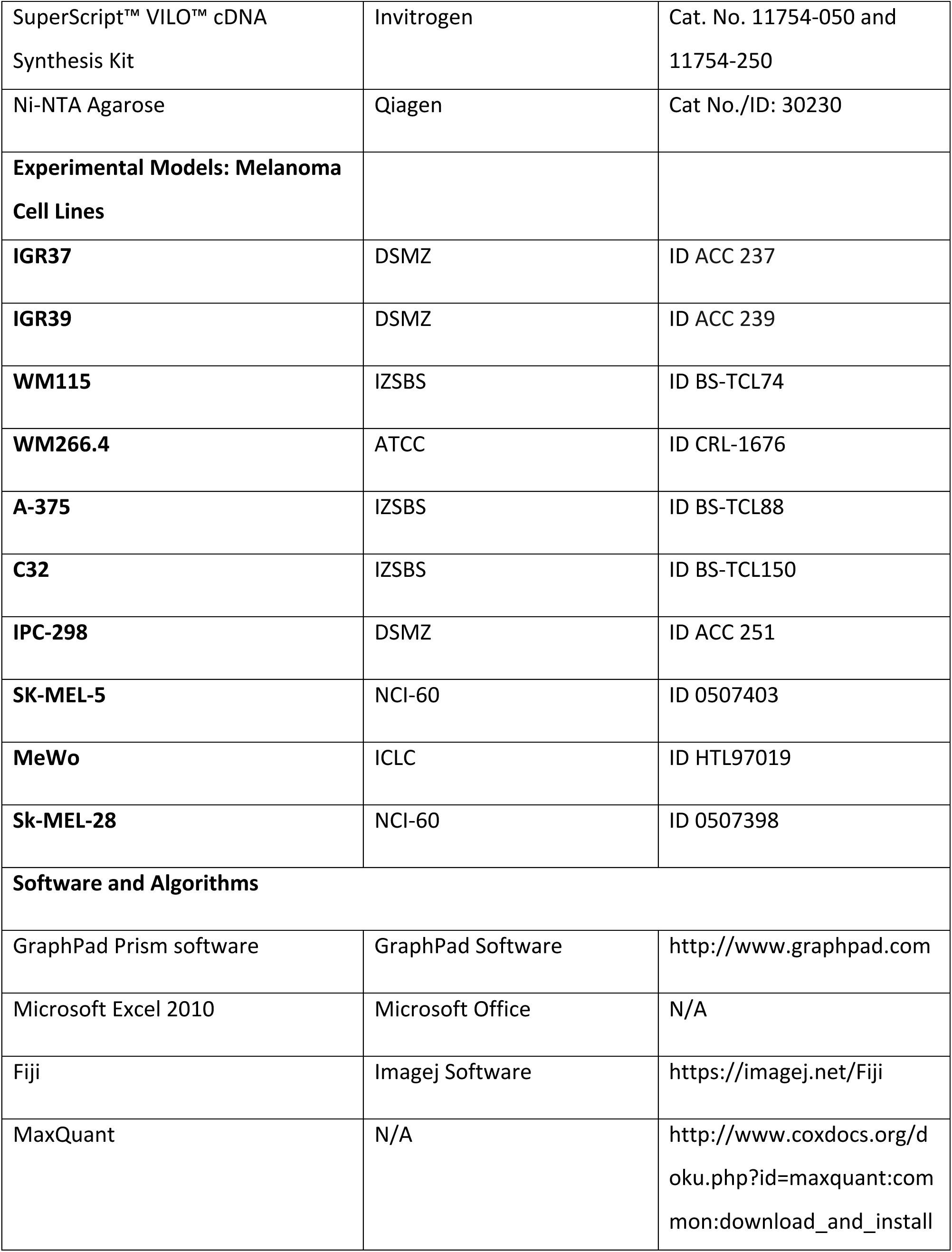
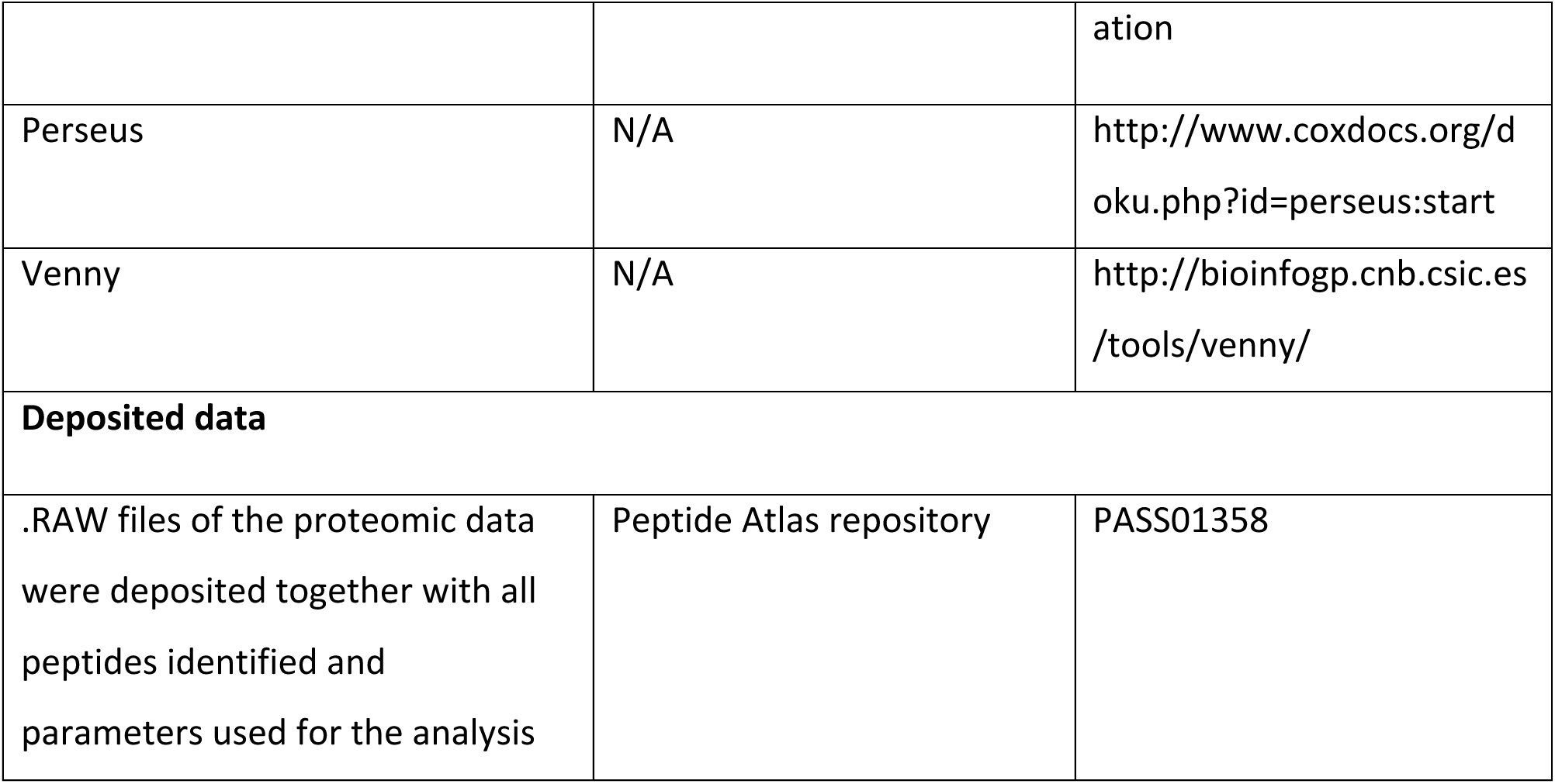

## Methods and Protocols

### Cell Culture

Human melanoma cell lines such as IGR37, IGR39 and IPC-298 were purchased from DSMZ; MEWO cell line was purchased from ICLC; SK-MEL-5 and SK-MEL-28 were purchased from NCI-60; WM266.4 was purchased from ATCC; WM115, A-375 and C32 were purchased from IZSBS.

All the cells were cultured in Dulbecco Modified Eagle’s Medium DMEM+ 10% FBS S.A.+ 2 mM L-Glutamine except for IPC-298 that was cultured in RPMI-1640+ 10% FBS S.A.+ 2 mM L-Glutamine. Cell lines were tested for mycoplasma by mycoplasma PCR Test Kit.

### Analysis of human biopsies

Formalin-fixed paraffin-embedded tissues were sliced into serial 8-µm-thick sections and collected for immunohistochemical (IHC) staining. Human paraffin samples were stained for Haematoxylin/Eosin (Diapath) to assess histological features, according to standard protocol. For Ki67 immunoanalysis paraffin was removed with xylene and sections were rehydrated in graded alcohol. Tissue slides were incubated in 10% peroxidase solution for 1 h at 65 °C to remove melanin pigments and then antigen retrieval was carried out using preheated target retrieval solution for 45 min at 95 °C. Tissue sections were blocked with FBS serum in PBS for 60 min and incubated overnight with primary antibody (Thermoscientific, 1:50). The antibody binding was detected using a polymer detection kit (GAR-HRP, Microtech) followed by a diaminobenzidine chromogen reaction (Peroxidase substrate kit, DAB, SK-4100; Vector Lab). All sections were counterstained with Mayer’s hematoxylin, mounted in Eukitt (Bio-Optica) and then visualized with an Olympus BX51 or an Olympus BX63 upright widefield microscope using NIS-Elements (Nikon, Tokyo, Japan) or MetaMorph 7.8 software (Molecular Devices, San Jose, CA, USA), respectively. For Proteostat aggresome detection deparaffinized and rehydrated slides were fixed in 4% PFA for 15 min, incubated in Proteostat solution (1:2000, Proteostat Aggresome Detection Kit, Enzo) for 3 min and then destained in 1% Acetic acid for 20 min at room temperature. To visualize the cell nuclei, human slides were counterstained with 4,6-diamidino-2-phenylindole (DAPI, Sigma-Aldrich), mounted with a Phosphate-Buffered Salines/glycerol solution and examined with confocal or widefield microscopy. Confocal microscopy was performed on a Leica TCS SP5 confocal laser scanning based on a Leica DMI 6000B inverted motorized microscope. The images were acquired with a HC FLUOTAR L 25X/NA0.95 VISIR water immersion objective using the 405 nm and the 488 nm laser lines. The software used for all acquisitions was Leica LAS AF. Widefield microscopy was performed on an Olympus BX63 upright microscope equipped with a motorized stage for mosaic acquisitions and with both Hamamatsu ORCA-AG black and white camera and Leica DFC450C color camera. The mosaic images were acquired using a UPlanSApo 4X/NA0.16 dry objective with MetaMorph 7.8 software (Molecular Devices). Quantitative analysis of stained signals was performed using ImageJ software. Both protein aggregates (dots) and nuclei were analyzed. Dots were normalized on nuclei and T-test statistical analysis was performed to estimate the differences between primary (N=6) and metastatic tissues (N=6). All the analysis was done in technical duplicates.

### Time lapse microscopy

For live cell imaging experiments, melanoma cells were cultured in six-well plates (5×10^3^ cells per plate). Cultures were transferred to a live cell imaging workstation composed by an Olympus IX81 inverted microscope equipped with motorized stage and a Hamamatsu ORCA-Flash4.0 camera. The images were collected every 5 min for a total recording time of 72 h for each dish using a LUC Plan FLN 20X/NA0.45 Ph dry objective with CellSens software (Olympus). The analysis was done, in biological triplicate, by using trackmate (Fiji).

### Secretome preparation from cell cultures and SILAC labeling

All melanoma cells were grown in a DMEM except for IPC-298 which was grown in a RPMI medium, complemented with essential amino acids Arg and Lys, containing naturally occurring atoms (Sigma) (the light medium) or two of their stable isotope counterparts (the medium and heavy media) (Cambridge Isotope Laboratories, Inc.; CIL). The medium culture contained arginine (L-Arg 13C6-14N4) and lysine (L-Lys 13C6-15N2) and the heavy culture contained arginine (L-Arg 13C6-15N4) and lysine (L-Lys 13C6-15N2) amino acids. After five cell divisions to obtain full incorporation of the labeled amino acids into the proteome, cells were counted and equal numbers of cells were split to 15 cm dishes at roughly 50% confluence. Once cell lines reached ∼70% confluence, one 15-cm dishes of each cell line were washed 3x with PBS and 3x with serum-free media. Cells were starved in serum free media for 18 h, the conditioned media (CM) was centrifuged (2000 rpm, 3 min), filtered (0.22 µM) to remove detached cells and concentrated via centrifugation at 6 000 RPM in 10 kDa molecular weight cutoff concentrating columns. Then, 500 µl of concentrated medium was filtered by microcon filters with 10 kDa cutoff (Millipore) and buffer was exchanged with Urea 8 M Tris100 mM or PBS.

### Protein aggregates detection

Aggregates Flourescence measurement was performed following manufacturer’s instructions (http://www.enzolifesciences.com/fileadmin/files/manual/ENZ-51023_insert.pdf). Briefly, 2 µL of the diluted PROTEOSTAT®Detection Reagent were added into the bottom of each well of a 96 well microplate. 98 µL of the secreted proteins in PBS were added to each well. Protein concentration was 10 µg/mL. The final concentration of the PROTEOSTAT detection dye was 1000 fold dilution in the assay. We run control samples as well as 1X Assay Buffer alone (no protein), as a blank sample. The microplate containing samples was incubated in the dark for 15 min at room temperature. Generated signals were read with a fluorescence Microplate reader (Tecan Infinite 200) using an excitation setting of about 550 nm and an emission filter of about 600 nm.

### Secretome analysis

Proteins secreted by 2×10^6^ cells were replaced in Urea 8 M Tris100 mM pH 8 and sonicated with BIORUPTOR (3 cycles: 30 seconds on/ 30 seconds off). By using microcon filters with 10 kDa cutoff (Millipore), cysteines reduction and alkylation was performed adding TCEP (Thermo scientific) 10 mM and 2-Chloroacetamide (Sigma-Aldrich) 40 mM in Urea 8 M Tris100 mM pH 8 for 30 min at room temperature; as described for the FASP protocol. Buffer was exchanged by centrifugation at 10000 rpm for 10 min and PNGase F (New England Biolabs) (1:100= enzyme: secreted proteins) was added for 1 h at room temperature following manufacturer’s instruction. Buffer was again exchanged by centrifugation at 10000 rpm for 10 min with ammonium bicarbonate 50 mM and proteins were in solution digested with trypsin (Trypsin, Sequencing Grade, modified from ROCHE) (1:50= enzyme: secreted proteins) overnight at 37 °C. Peptides were recovered on the bottom of the microcon filters by centrifugation at 10000 rpm for 10 min and on the top, adding two consecutive wash of 50 µl of NaCl 0.5 M. The undigested polypeptides on the top of the filters were further digested with GluC (Endoproteinase Glu-C Sequencing Grade ROCHE) (1:50= enzyme: secreted proteins) overnight at 37 °C upon buffer exchange with phosphate buffer (pH 7.8). Eluted peptides were purified on a C18 StageTip. 1 µg of digested sample was injected onto a quadrupole Orbitrap Q-exactive HF mass spectrometer (Thermo Scientific). Peptides separation was achieved on a linear gradient from 95% solvent A (2% ACN, 0.1% formic acid) to 55% solvent B (80% acetonitrile, 0.1% formic acid) over 75 min and from 55% to 100% solvent B in 3 min at a constant flow rate of 0.25 µl/min on UHPLC Easy-nLC 1000 (Thermo Scientific) where the LC system was connected to a 23-cm fused-silica emitter of 75 µm inner diameter (New Objective, Inc. Woburn, MA, USA), packed in-house with ReproSil-Pur C18-AQ 1.9 µm beads (Dr Maisch Gmbh, Ammerbuch, Germany) using a high-pressure bomb loader (Proxeon, Odense, Denmark).

The mass spectrometer was operated in DDA mode as described previously (Matafora et al, 2017): dynamic exclusion enabled (exclusion duration = 15 seconds), MS1 resolution = 70,000, MS1 automatic gain control target = 3 x 106, MS1 maximum fill time = 60 ms, MS2 resolution = 17,500, MS2 automatic gain control target = 1 x 105, MS2 maximum fill time = 60 ms, and MS2 normalized collision energy = 25. For each cycle, one full MS1 scan range = 300-1650 m/z, was followed by 12 MS2 scans using an isolation window of 2.0 m/z.

All the proteomic data as raw files, total proteins and peptides identified with relative intensities and search parameters have been loaded into Peptide Atlas repository (**PASS01358**).

### MS analysis and database search

MS analysis was performed as reported previously (Matafora et al, 2015). Raw MS files were processed with MaxQuant software (1.5.2.8), making use of the Andromeda search engine (Cox et al, 2011). MS/MS peak lists were searched against the UniProtKB Human complete proteome database (uniprot_cp_human_2015_03) in which trypsin and GluC specificity was used with up to two missed cleavages allowed. Searches were performed selecting alkylation of cysteine by carbamidomethylation as fixed modification, and oxidation of methionine, N-terminal acetylation and N-Deamination as variable modifications. Mass tolerance was set to 5 ppm and 10 ppm for parent and fragment ions, respectively. A reverse decoy database was generated within Andromeda and the False Discovery Rate (FDR) was set to <0.01 for peptide spectrum matches (PSMs). For identification, at least two peptides identifications per protein were required, of which at least one peptide had to be unique to the protein group.

### Quantification and Statistical Analysis

Silac and Label free from DDA .raw files were analyzed by MaxQuant software for protein quantitation and, depending from the experiment, SILAC Ratio or LFQ intensities were used. Statistical analysis was performed by using Perseus software (version 1.5.6.0) included in MaxQuant package. T-test and ANOVA statistical analysis was performed applying FDR<0.05 or P-value <0.05 as reported. KEGG enrichment pathway analysis was performed via EnrichR (http://amp.pharm.mssm.edu/Enrichr), using the Gene ID of the identified proteins.

### Oxysterol Quantification

Oxysterols were prepared from melanoma cell lines using a modified version of the protocol described by Griffiths et al. (Griffiths et al, 2013), consisting in an alcoholic extraction and a double round of reverse-phase (RP) solid-phase extraction (SPE) (Soncini et al, 2016). Briefly, melanoma cell pellets (2×10^6^ cells) were sonicated for 5 min by adding 1.0 ml of ethanol supplemented with + 20 pmol of each deuterated standard. 400 µl H_2_O were added and sonicated for other 5 min (final volume 1.5 ml of 70% ethanol). Upon centrifugation at volume finale 1.5 ml di 70% ethanol, supernatant was collected. The extract was applied to a preconditioned Sep-Pak tC18 cartridge (Waters). The oxysterol-containing flow-through was collected, together with the first 70% (vol/vol) ethanol wash. The collected oxysterols were vacuum-evaporated and reconstituted in 100% (vol/vol) isopropanol, diluted in 50 mM phosphate buffer, and oxidized by cholesterol oxidase addition. The reaction was stopped by methanol. Reactive oxysterols were then derivatized by Girard P reagent (TCI Chemical) and further purified by reverse phase chromatography using a Sep-Pak tC18 cartridge to eliminate the excess of GirardP reagent. Purified oxysterols were diluted in 60% (vol/vol) methanol and 0.1% formic acid. Eight µl of sample was resolved by on a nano-HPLC system connected to a 15-cm fused-silica emitter of 75 µm inner diameter (New Objective, Inc. Woburn, MA, USA), packed in-house with ReproSil-Pur C18-AQ 1.9 µm beads (Dr Maisch Gmbh, Ammerbuch, Germany) using a high-pressure bomb loader (Proxeon, Odense, Denmark). It was used a 12-min gradient from 20% to 100% of solvent B [63.3% (vol/vol) methanol, 31.7% (vol/vol) acetonitrile, and 0.1% formic acid], where solvent A is composed of 33.3% methanol, 16.7% acetonitrile, and 0.1% formic acid. Eluting oxysterols were acquired on a quadrupole Orbitrap Q-Exactive HF mass spectrometer (Thermo Scientific), where the survey spectrum was recorded at high resolution (R = 140,000 at 200 m/z) and the five most intense peaks were further fragmented. The identification of the oxysterols species was made by comparing the retention times of the analytes with those of the synthetic, deuterated standards previously run on the same system in the same chromatographic conditions. The quantification was achieved by means of stable-isotope dilution MS using internal standards. The total ion current for derivatized oxysterols was extracted for each acquisition, areas of the peaks were integrated manually using Xcalibur software, and the absolute amount of oxysterols was determined by comparing their areas with those of the internal standards, using the following equation:

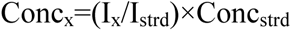

### Protein quantification

Protein quantification was performed using Bradford assay (Bio-Rad). For each sample, the absorbance was measured by a spectrophotometer at a wavelength of 595 nm. Sample protein concentration was determined based on a Bovine Serum Albumin (BSA) standard curve.

### Western Blot Assays

For western blot analyses, proteins were extracted in buffer containing Urea 8 M, TrisHCl 100 mM pH 8. Briefly, cell lysates (50 µg) were separated by SDS-PAGE using a precast polyacrylamide gel with a 4% to 12% gradient (Invitrogen). After the electrophoretic run, proteins were transferred onto a 0.22 μm nitrocellulose membrane (Amersham Protran, GE Healthcare) in wet conditions. The assembled sandwich was loaded in a Trans-Blot Cell (Bio-Rad) and immersed in 1X cold Tris-Glycine transfer buffer with the addition of 20% methanol. The transfer was allowed over-night at constant voltage (30 V). Correct protein transfer was verified staining the membrane with Ponceau red (Sigma Aldrich) for few seconds. After washing the membrane with Tris-buffered Saline-Tween 20 (TBST, 1X TBS with 0.1% Tween-20) non-specific binding of antibodies was blocked by adding 5% low-fat dry milk in TBST for 1 hour at room temperature. Murine Anti-human Pmel17 (HMB45 Thermo scientific) primary monoclonal antibody, was diluted in the same blocking solution to a final concentration of 1:100. The anti-HDAC2 antibody (Cell-Signalling) was used to normalize the amount of proteins loaded onto the gel. Anti-murine IgG1 secondary antibody conjugated with the enzyme horseradish peroxidase (HRP) was used to a final concentration of 1:2000 in 5% milk-TBST.

### Immunofluorescence

Cells were fixed and permeabilized as describe previously (Matafora et al, 2009). After treatment, cells were fixed with 4% (wt/vol) paraformaldehyde, blocked with PBS-BSA (1% wt/vol), made permeable with Triton X-100 0.2% (Sigma-Aldrich) for 3 min, and incubated with proteostat (1:1000) or specific antibodies diluted in 0.2% bovine serum albumin in PBS. Cells were then washed three times with PBS and stained with DAPI (Sigma-Aldrich). Cell were observed by confocal microscopy performed on a Leica TCS SP5 or a Leica TCS SP2 AOBS confocal laser scanning. The confocal systems were respectively based on a Leica DMI 6000B or a DM IRE2 inverted microscope equipped with motorized stage. The images were acquired with an HCX PL APO 63X/NA1.4 oil immersion objective using the 405 nm, 488 nm or 561 nm laser lines. The software used for all acquisitions was Leica LAS AF (on TCS SP5 system) or Leica Confocal Software (on TCS SP2 AOBS System).

### Recombinant PMEL (rMα) expression and purification

The luminal fragment of PMEL, rMα, consisting of amino acids 25–467 was subcloned from PGEX vector into a pET28a vector, in order to have 6xHis tag at the N-terminus, and expressed in BL21-DE3 E. Coli. Shaken cultures were grown at 37 °C to OD600=0.5 in the presence of kanamycin and then induced with 1 mM IPTG for 4 h. Cells were collected via centrifugation at 4 °C, resuspended in TBS (Tris-Buffered Saline: 150 mM NaCl, 50 mM Tris-HCl, pH 7.6), and frozen at −80 °C. The resuspended pellet was thawed and the cells were lysed by probe sonication. rMα formed inclusion bodies that were collected by centrifugation after three washings in 1,5M NaCl, 100mM Tris-HCl pH 7.4, 1% Triton-X 100 buffer and two in TBS (Fowler et al, 2006). The inclusion bodies were dissolved in 9M Urea, 100 mM NaH_2_PO_4_, 10 mM Tris-HCl pH 8.0 and then filtered through a 0.22 *μ*M cellulose acetate filter and stored at room temperature. The protein was purified using Ni-NTA agarose beads (Qiagen, Germany) under denaturing condition. Briefly, 2 ml 50% slurry of Ni-NTA agarose beads were equilibrated with binding buffer (9M Urea, 100mM NaH_2_PO_4_, 10mM Tris-HCl pH 8.0) before adding the sample. After binding for 1h and 30 min, two washes were performed with 9M Urea, 100mM NaH_2_PO_4_, 10mM Tris-HCl pH 6.5; elution was obtained in 9M Urea, 100mM NaH_2_PO_4_, 10mM Tris-HCl pH 4.5.

### PMEL aggregates refolding and administration to cells

Recombinant PMEL aggregates refolding was obtained by slightly modifying a previously described protocol (Fowler et al, 2006). In particular, we did sequential dilutions form denaturing to native condition by performing first gel filtration and then buffer exchange. Briefly, after NiNTA purification, recombinant PMEL was subjected to gel filtration in mild denaturating buffer (4M Urea, 100 mM Tris-HCl pH 8.0) on a Superdex 200 16/60 column (GE Healthcare Life Sciences, USA), in order to allow partial refolding and avoid the elution of the protein in the void volume. The fractions corresponding to PMEL elution were pulled together and concentrated by using Amicon Ultra centrifugal tubes with 10 kDa cutoff (Millipore, USA). To allow a complete refolding and cells culture treatment the buffer was exchanged with PBS. Recombinant PMEL aggregates were administered to cells in culture media at a final concentration of 0,5 μM.

### IGR39 treatment with IGR37 conditioned medium

IGR39 were seeded at the concentration of 100000 cells/well. At the same time IGR37 were seeded at 60% confluency in a 10 cm petri dish. After 24 h the media deriving from IGR37 were filtered on a 0.22 μM cellulose acetate filter, in order to remove dead cells and cells debris, and administered to IGR39. After 24 h, cells were harvested and RNA extraction was performed.

### RNA extraction, RT-PCR and real-time PCR

Total RNA was extracted using Maxwell RSC simply RNA (Promega, USA) according to manufacturer’s instructions, and RNA was quantified by nanodrop. 1μg of total RNA was used for retro-transcription using SuperScript™ VILO™ cDNA Synthesis Kit (Invitrogen, USA). cDNA was diluted 1:10 and qPCR were performed using LightCycler^®^ 480 SYBR Green I Master (Roche, Switzerland). The primer sequences are provided in Supplementary Table S11. Expression data were normalized to the geometric mean of the housekeeping gene RPLP0 to control the variability in expression levels and were analyzed using the 2-ΔΔCT method. Primers for qPCR: CTGF-Forward primer: GGGAAATGCTGCGAGGAGT, CTGF-Reverse primer: GCCAAACGTGTCTTCCAGTC; RPLP0-Forward primer: GTTGCTGGCCA ATAAGGTG, RPLP0-Reverse primer: GGGCTGGCACAGTGACTT.

### Cell viability assays

Melanoma cell lines were seeded into 6-well plates. MUSE reagent was added to detached cells and cell viability was assessed according to the manufacturer’s instructions (http://www.merckmillipore.com/IT/it/product/Muse-Count-Viability-Assay-Kit-100-Tests,MM_NF-MCH100102#anchor_UG). Viability was accessed by measuring cell confluence (%) and number of dead and alive cells by using Muse™ Cell Analyzer.

### MTT cell viability assay

To perform 3-(4,5-dimethylthiazol-2-yl)-2,5-diphenyltetrazolium bromide (MTT; Sigma) cell viability assay, melanoma cells were seeded in 96-well plates (5×10^3^ cells/well) and were treated with doxorubicin or/and NB360 as indicated in the text. At the end of the experiments, the cell cultures were supplemented with 150 µl of 0.5 mg/ml MTT assay and incubated for an additional 4 h. Then, equal volume of solubilizing solution (dimethyl sulfoxide 40%, SDS 10% and acetic acid 2%) was added to the cell culture to dissolve the formazan crystals and incubated for 10 min at room temperature. The absorbance rate of the cell culture was detected at 570 nm by using a Microplate Reader (Bio-Rad, Hercules, CA, USA). Each experiment was performed as biological quadruplicate.

### Clonogenic assay

Melanoma cells (2000 cells/well) were seeded into 6-well plates and following cell attachment they were treated with DMSO or NB360 as indicated. Then, the plates were incubated at 37 °C with 5% CO_2_, until cells formed colonies (12–15 days). Colonies were fixed with 75% methanol and stained with 0.5% crystal violet, then rinsed with PBS, dried and counted using the ImageJ software.

## Acknowledgments

We thank the IFOM Functional Proteomics group and the Proteomics facility for critical comments and suggestions. We thank Luca Azzolin and Stefano Piccolo for YAP antibody and for their comments on our work. We acknowledge the Imaging, Mass Spectrometry and the Histopathology units at IFOM for their precious work. We thank Giannino Del Sal for discussion. We acknowledge Giuseppe Ossolengo for technical advice for gel filtration. We thank Jeffry W. Kelly for providing PMEL amyloidogenic fragment plasmid. Francesco Farris is a PhD student within the European School of Molecular Medicine (SEMM). We also thank Dr. Ulf Neumann from Novartis for kindly providing NB360 according to MTA agreement.

## Author Contributions

Methods development, V.M., U.R., and G.M.; Validation and Formal Analysis, V.M.; Investigation, V.M., G.M., F.F., U.R., S.T., C.B., F.P., and F.C.; Resources, A.B., E.B., S.M., and L.L.; Writing original draft, V.M., G.M., and A.B.; Supervision, Project Administration and Funding Acquisition, A.B.

## Study approval

Informed consent was obtained from all study participants. Study approval was given by the Institutional Review Board of the Grande Ospedale Metropolitano Niguarda. All cases of melanoma cancer were pathologically confirmed.

## Data availability

Proteomic datasets produced in this study are available in the following databases: Proteomics Identification database PeptideAtlas http://www.peptideatlas.org/

## Supplemental Information

**Table S1.** Proteins statistically significant in WM cell lines secretome (SILAC)

**Table S2.** Proteins statistically significant in IGR cell lines secretome (SILAC)

**Table S3:** Proteins statistically significant in the secretome of a cohort of melanoma cell lines (normalized on cell number)

**Table S4:** Proteins statistically significant in the secretome of a cohort of melanoma cell lines (normalized on protein concentration)

**Table S5:** Proteins statistically significant in the secretome of IGR cell lines treated with NB360

**Video S1-S2:** Time-lapse microscopy of IGRs cells

**Supplemental Figures S1–S12**

