## supplementary figures for "Amyloid aggregates accumulate in melanoma metastasis driving YAP mediated tumor progression"

**Supplemental Information:**

**Table S1.** Proteins statistically significant in WM cell lines secretome (SILAC)

**Table S2.** Proteins statistically significant in IGR cell lines secretome (SILAC)

**Table S3:** Proteins statistically significant in the secretome of a cohort of melanoma cell lines (normalized on cell number)

**Table S4:** Proteins statistically significant in the secretome of a cohort of melanoma cell lines (normalized on protein concentration)

**Table S5:** Proteins statistically significant in the secretome of IGR cell lines treated with NB360

**Video S1-S2:** Time-lapse microscopy of IGRs cells

**Supplemental Figures S1–S12**

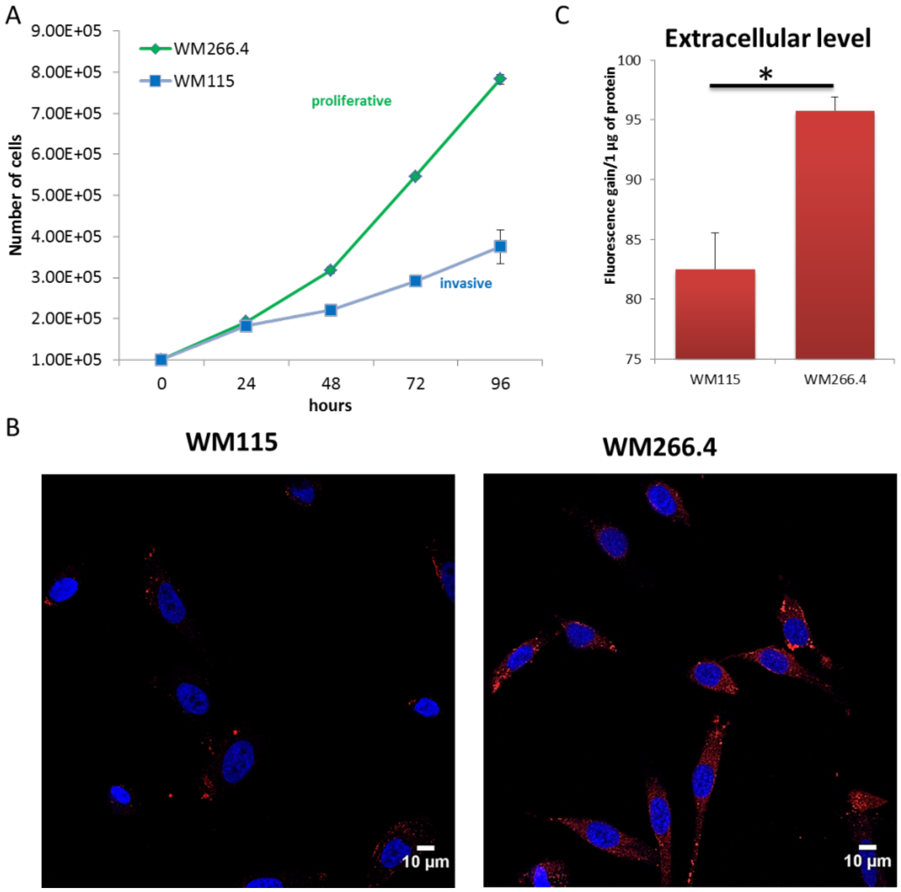


**Figure S1. Analysis of model system for primitive and metastatic melanoma.** (A) Growth curve of primitive WM115 and metastatic WM266.4 cell line (n=3). (B) Confocal immunofluorescence images of Proteostat (1:2000, red spots) and DAPI staining (blue) for WM155 and WM266.4 cell line. Scale Bar is 10µm. (C) Quantitation of aggregates in the extracellular space in WMs. Fluorescence gain of soluble proteins treated with Proteostat reagent. T-test analysis was performed, * = 0.01<p-value<0.05.

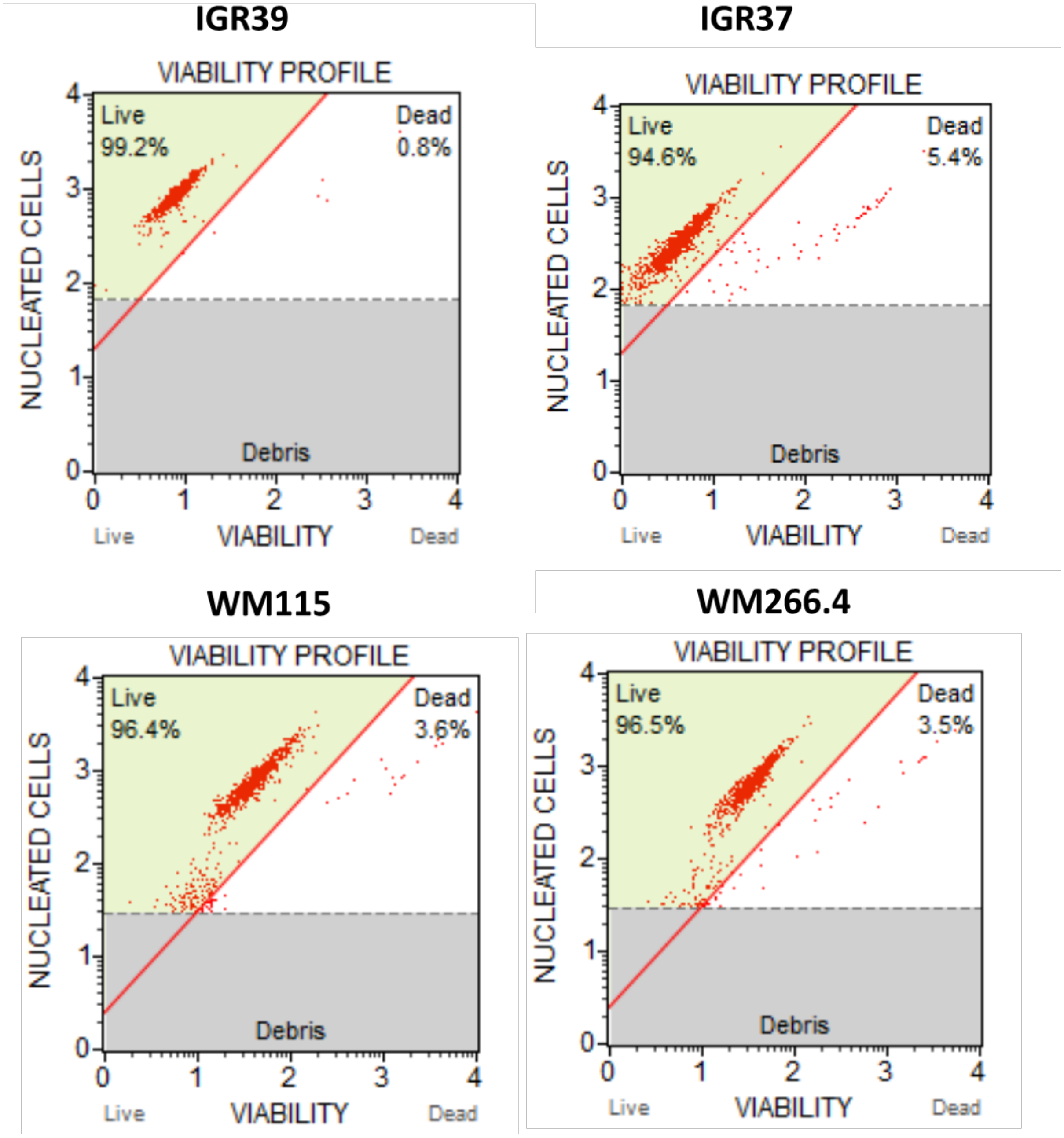


**Figure S2. Viability profile of IGRs and WMs cells.** After 24h starvation, viability was detected by measuring cell confluence (%) and number of dead and alive cells using Muse™ Cell Analyzer**.**

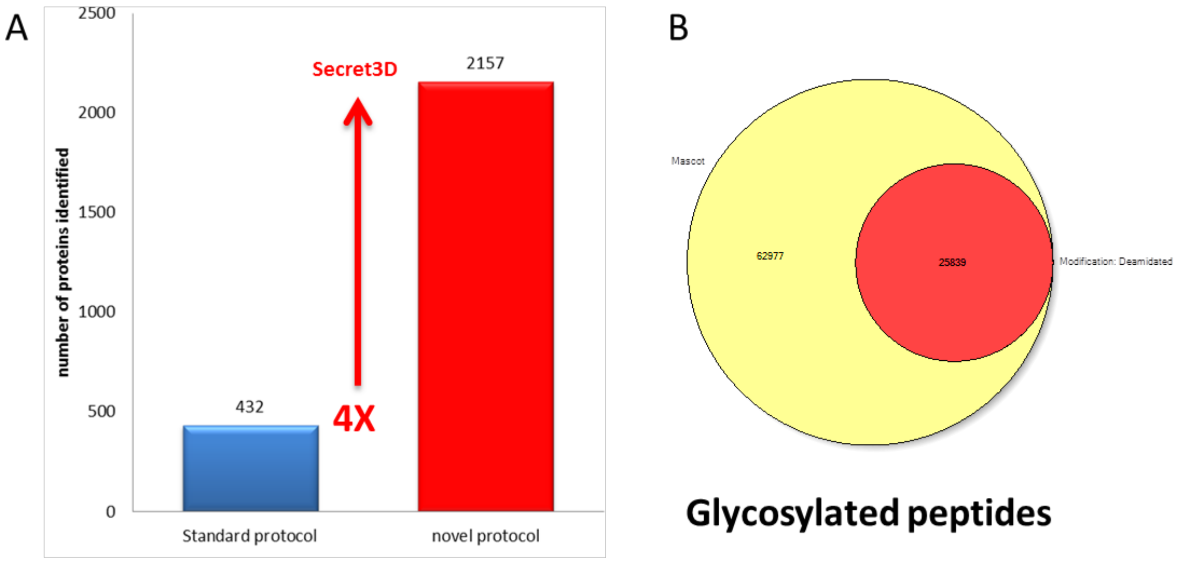


**Figure S3. Secretome analysis.** (A) Comparison of identified proteins by Secret3D or by standard protocol. (B) Number of total and glycosylated peptides identified by Secret3D method.

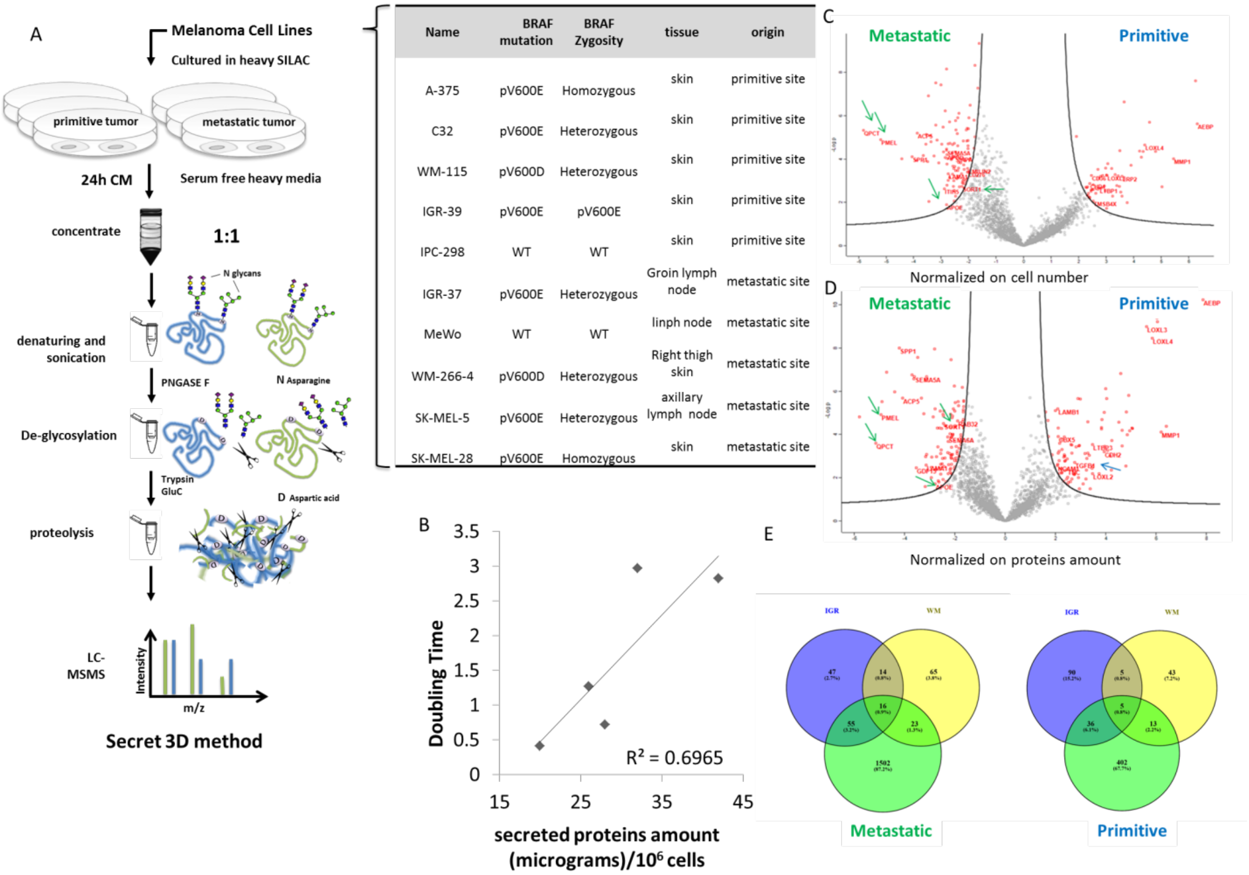


| Name | BRAF mutation | BRAF Zygosity | tissue | origin |
|---|---|---|---|---|
| A-375 | pV600E | Homozygous | skin | primitive site |
| C32 | pV600E | Heterozygous | skin | primitive site |
| WM-115 | pV600D | Heterozygous | skin | primitive site |
| IGR-39 | pV600E | pV600E | skin | primitive site |
| IPC-298 | WT | WT | skin | primitive site |
| IGR-37 | pV600E | Heterozygous | Groin lymph node | metastatic site |
| MeWo | WT | WT | linph node | metastatic site |
| WM-266-4 | pV600D | Heterozygous | Right thigh skin | metastatic site |
| SK-MEL-5 | pV600E | Heterozygous | axillary lymph node | metastatic site |
| SK-MEL-28 | pV600E | Homozygous | skin | metastatic site |

**Figure S4. Proteomic analysis of the secretome from primitive and metastatic melanoma cells**. A) MS workflow of Secret3D applied on melanoma cell lines described on the right. (B) Scatter plot of secreted proteins versus doubling time of the cell lines analyzed. (C, D) Volcano plots of the quantified proteins with both normalization strategies discussed in the main text. (E) Venn diagram of the significant proteins shared by all melanoma cell lines.

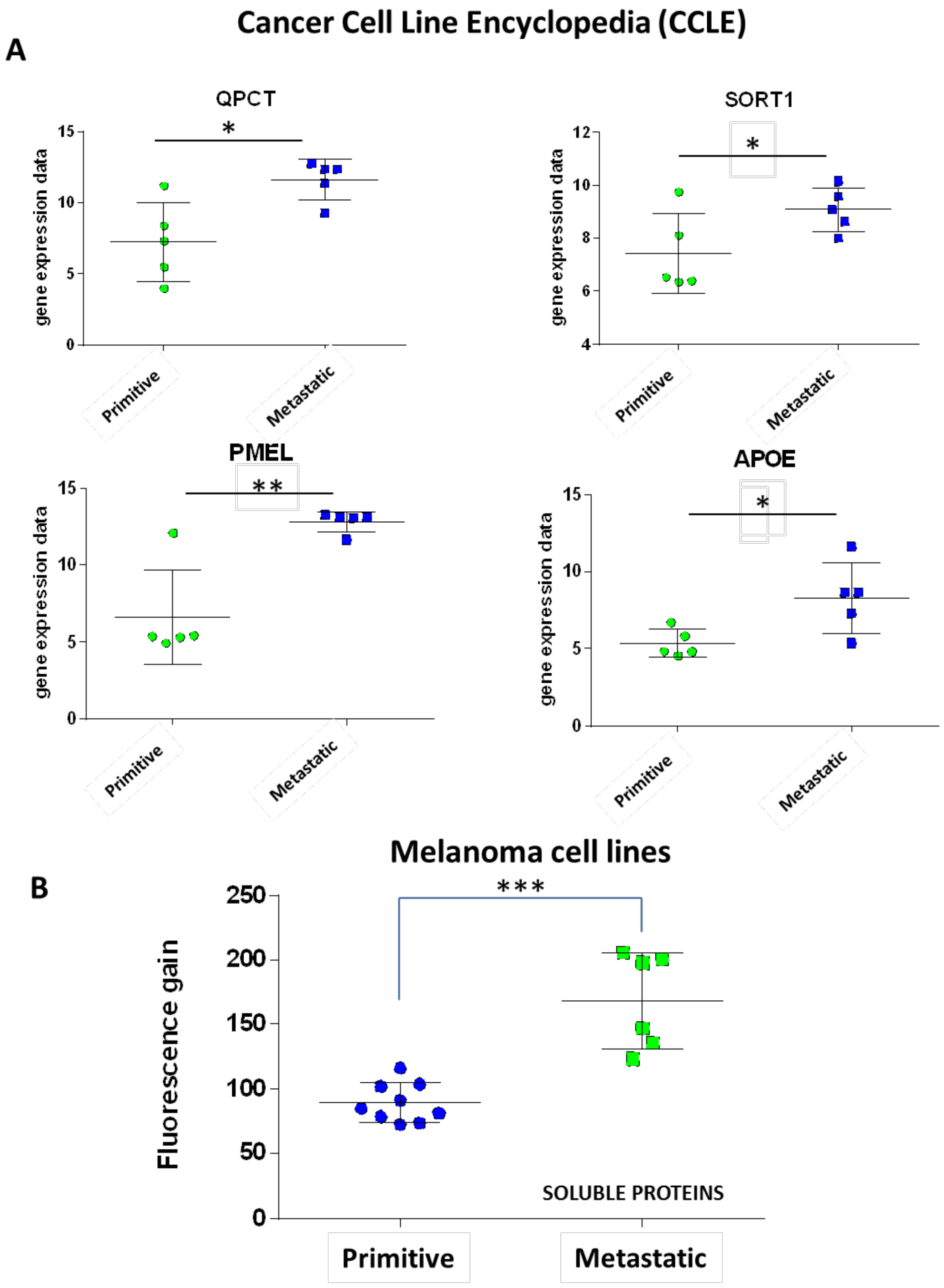


**Figure S5. PMEL and amyloidogenic proteins are enriched in metastatic cells secretome.** (A) Gene expression data from the Cancer Cell Line Encyclopedia (CCLE). (B) Fluorescence gain of soluble proteins treated with Proteostat reagent in the secretome of melanoma cells. T-test analysis was performed. * 0.01<p-value<0.05; ** 0.001<p-value<0.01; *** p-value <0.001.

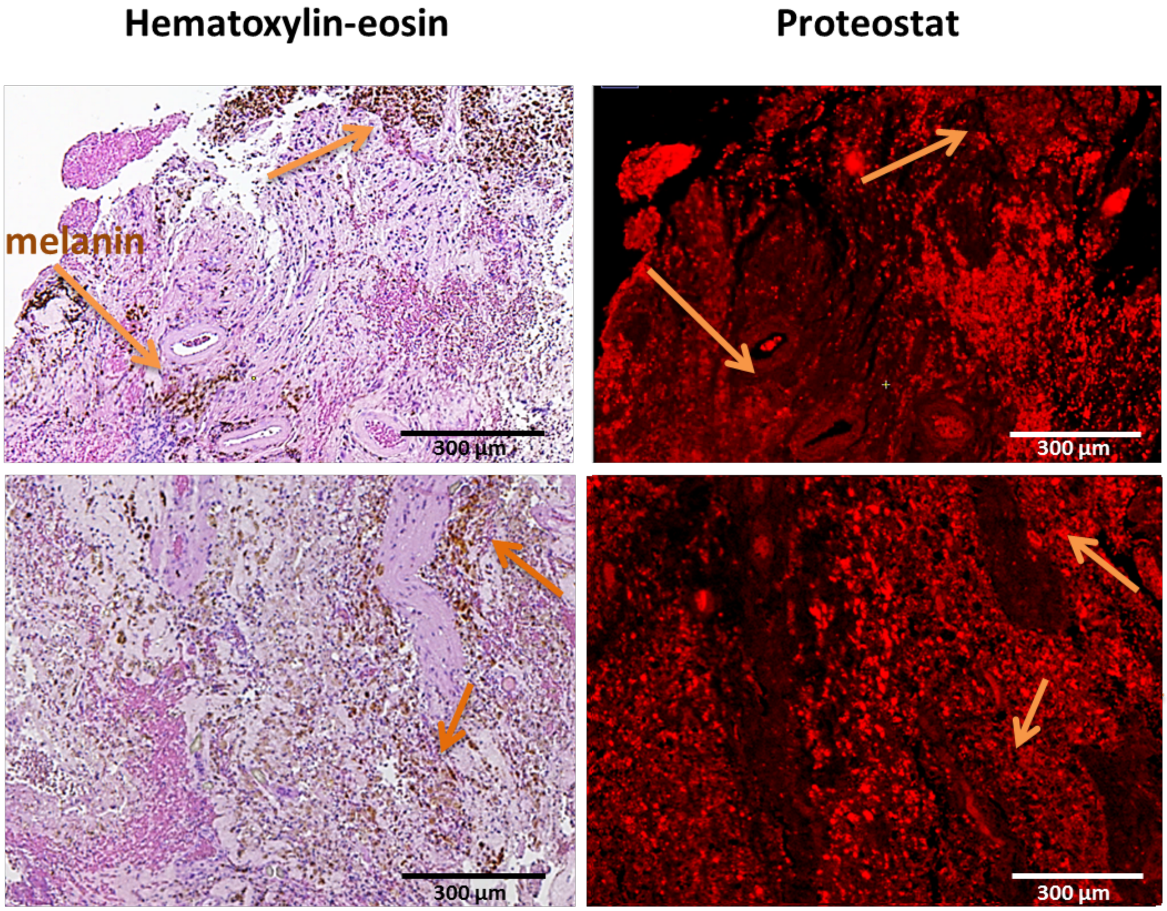


**Figure S6. Mosaic acquisitions of melanoma metastases in human brain stained with hematoxylin-eosin or with Proteostat.** The arrows point at melanin signals. Scale bar is 300µm.

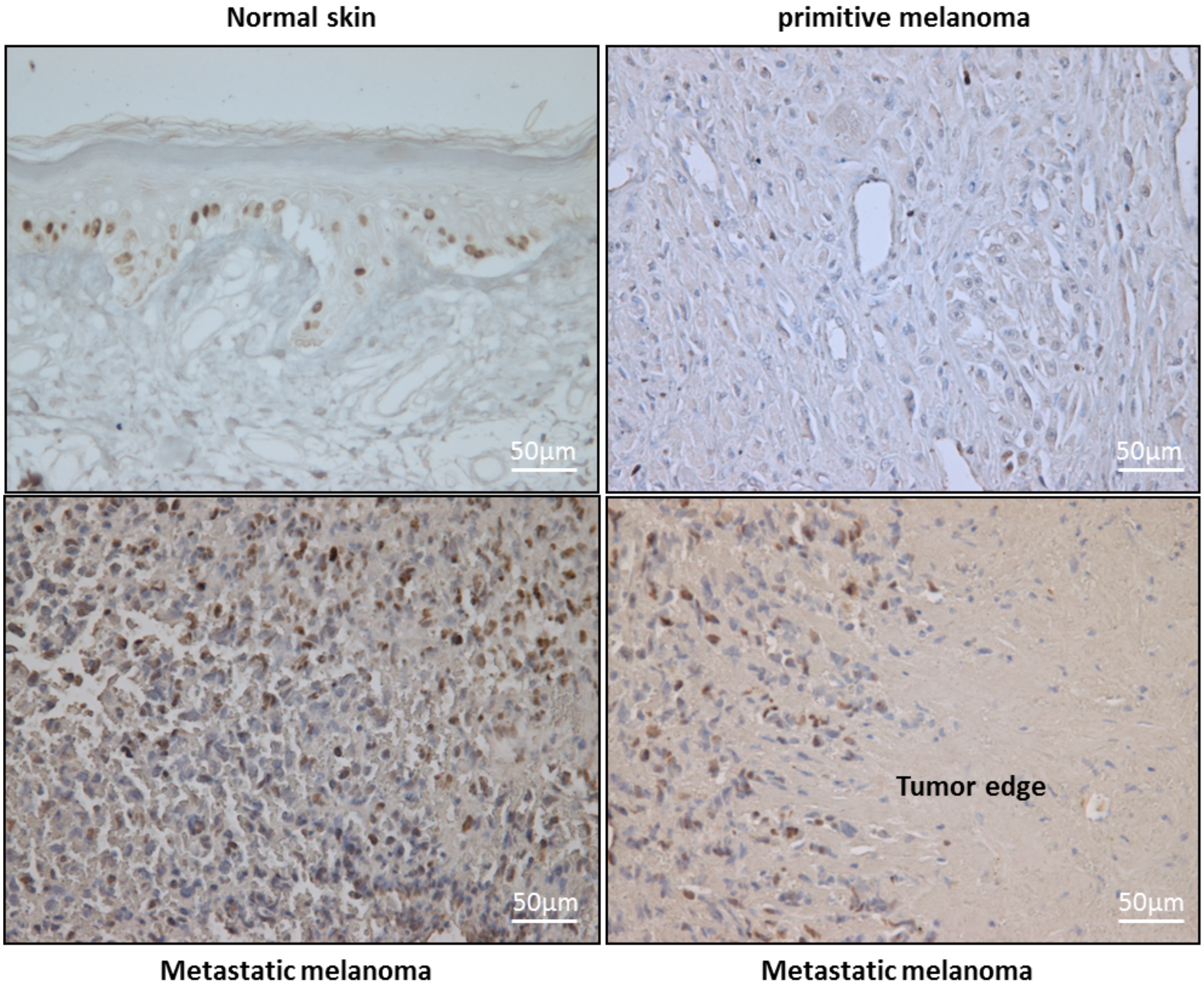


**Figure S7. KI-67 staining (brown) in healthy tissue, primitive melanoma and metastatic melanoma.** Scale bar is 50µm.

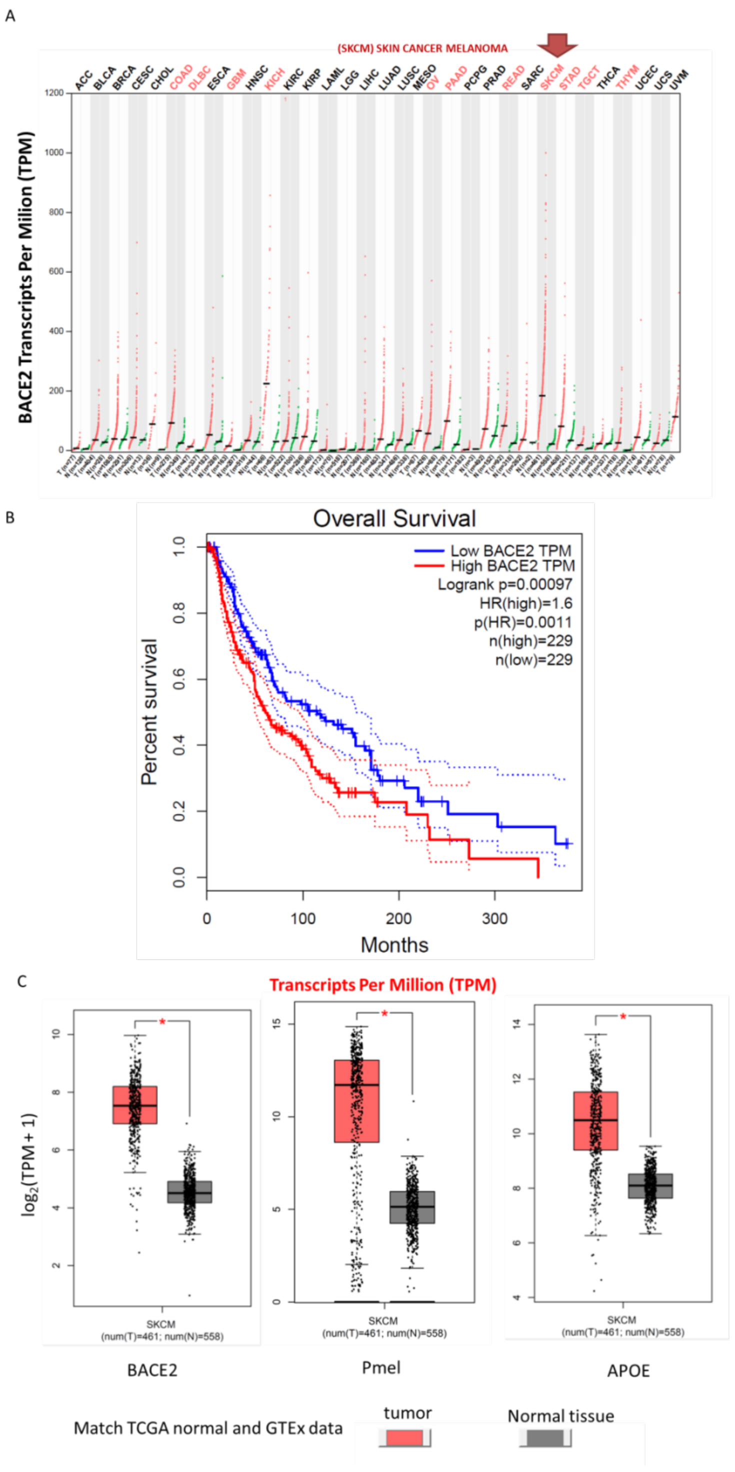


**Figure S8 Gene expression profiling in TCGA normal and GTEx dataset by using GEPIA software (http://gepia.cancer-pku.cn/).** (A) Survival analysis. (B) Gene expression by cancer type. (C) Gene expression cancer/normal comparison.

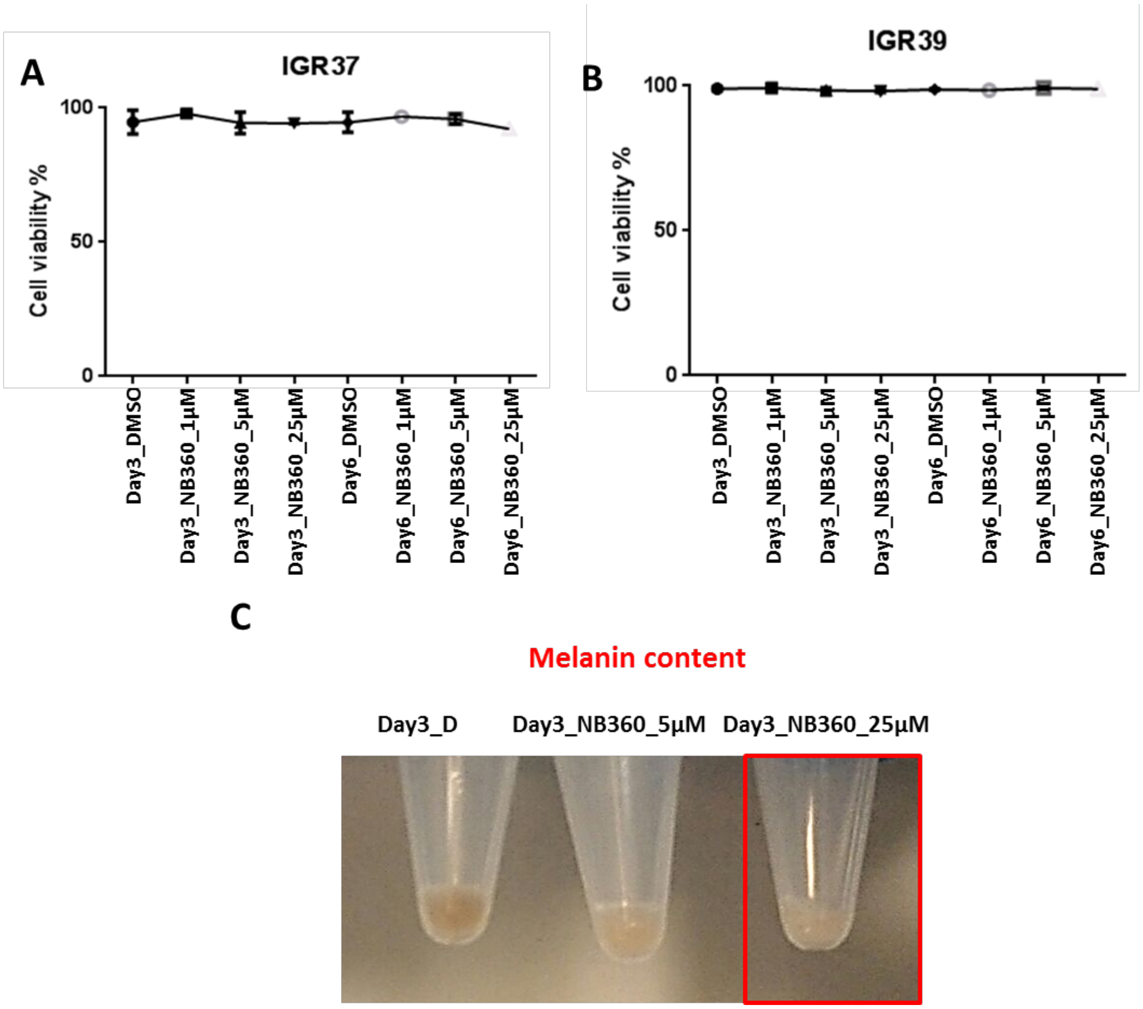


**Figure S9. Cell viability assay.** Cell viability of IGRs treated with NB-360 by using different concentrations and time of incubation as reported in the panels A and B. Pigmentation of IGR37 upon incubation with NB-360, panel C.

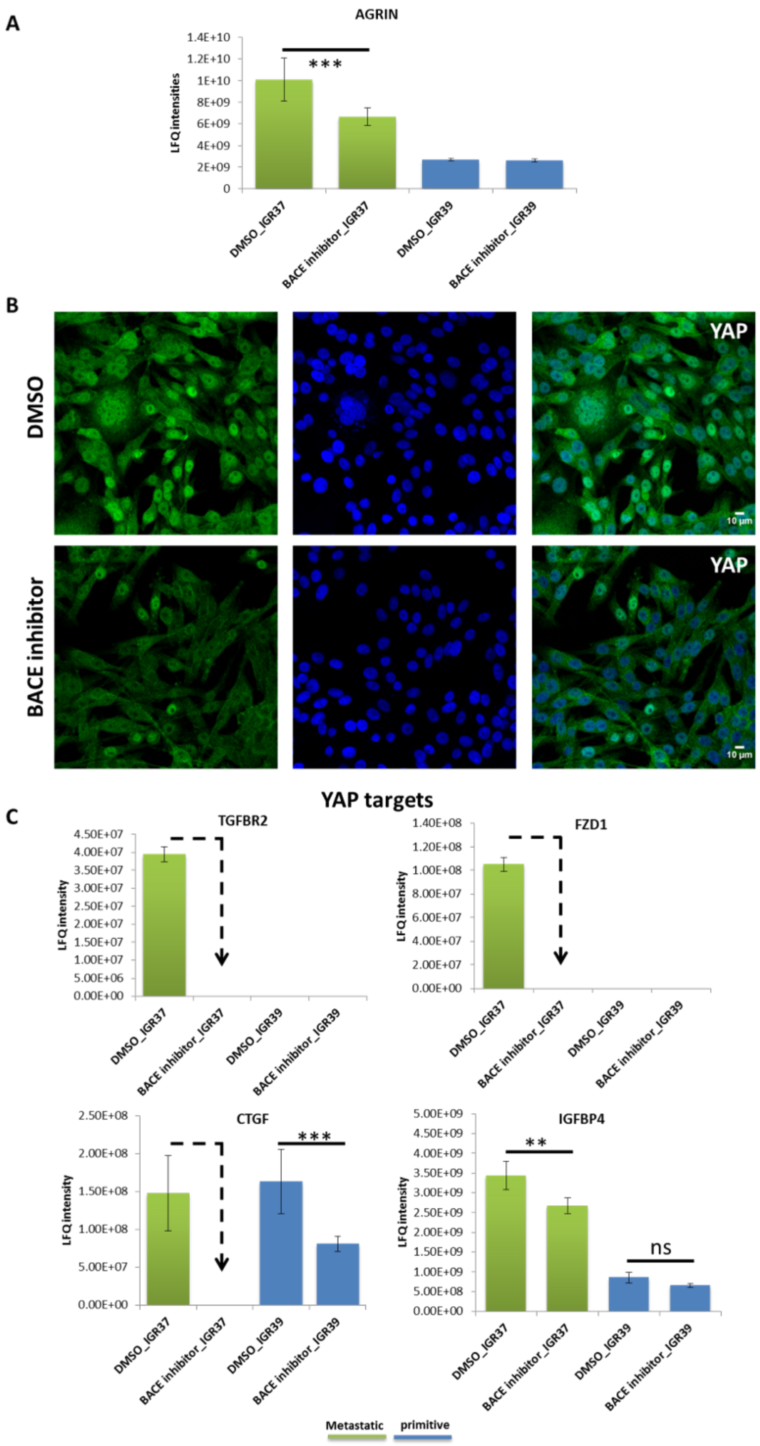


**Figure S10. NB360 affects YAP transcriptional activity. (A)** Label free intensities of agrin in IGRs treated with DMSO or BACE inhibitor. (B) Confocal fluorescence images of anti-YAP antibody signal (green) and DAPI staining (blue) in IGR 37 upon treatment with DMSO or BACE inhibitor. Scale Bar is 10µm. (C) Label free intensities of YAP targets in IGRs treated with DMSO or BACE inhibitor. T-test analysis was performed, ** = 0.001<p-value<0.01; *** = p-value <0.001.

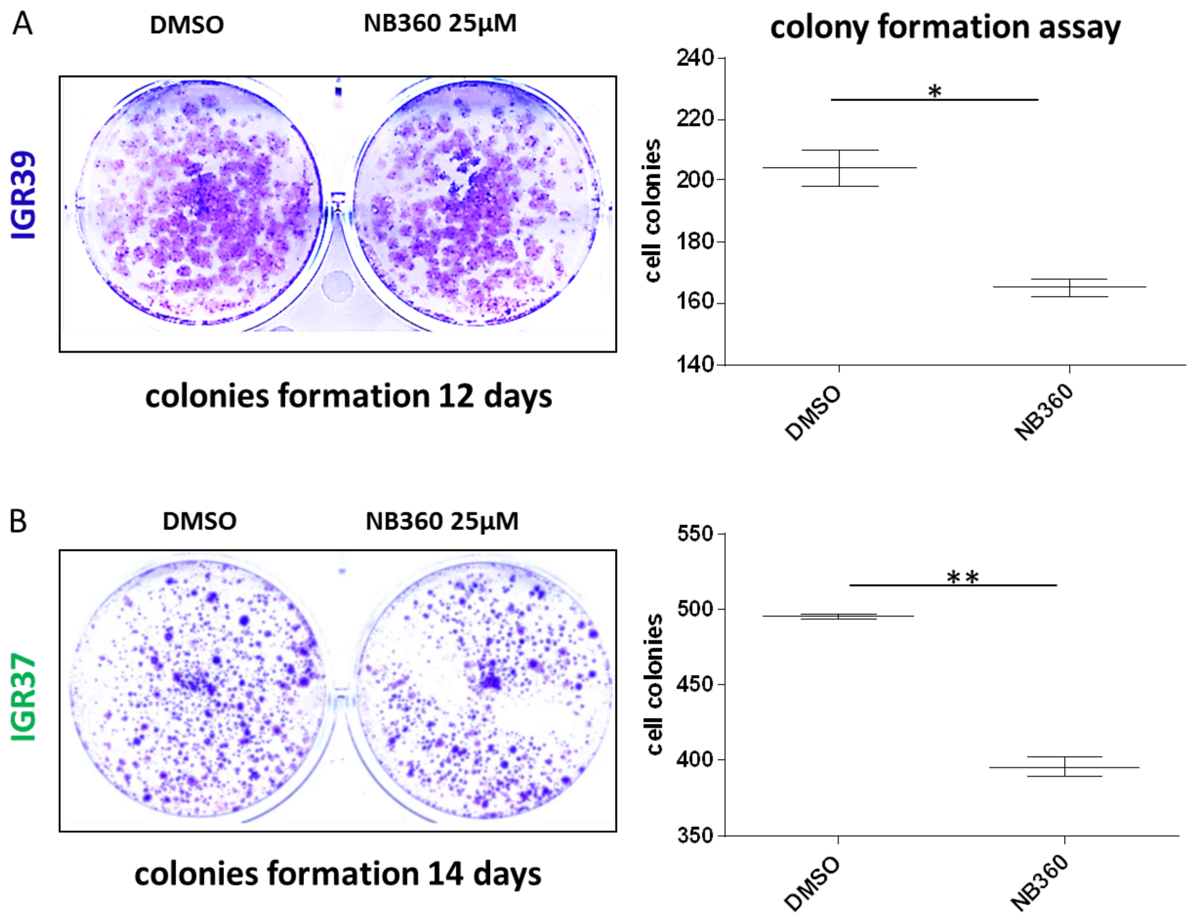


**Figure S11.** (A-B) Colony formation assay and relative quantitation on the right panels, for IGR39 and IGR37 treated with DMSO or NB-360. Biological replicates N=2. T-test analysis was performed. * = 0.01<p-value<0.05; ** = 0.001<p-value<0.01.

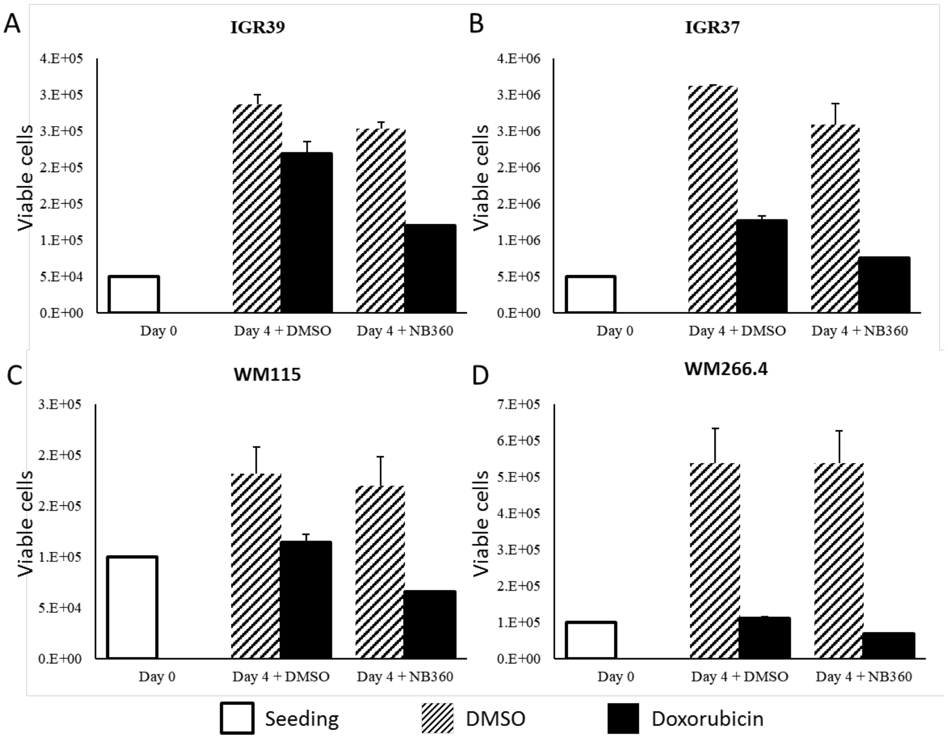


**Figure S12. Viability of WMs and IGRs melanoma cell lines upon doxorobucin and/ot NB360 treatment.** Cells were treated with DMSO, doxorubicin 10 µM, NB-360 25µM and the combination of doxorubicin plus NB-360. Viability was assessed by measuring cell confluence (%) and number of dead and alive cells using the Muse™ Cell Analyzer.
